# Benchmarking Large Language Models for Assay-Aware Drug-Target Binding Affinity Prediction

**DOI:** 10.64898/2026.10.11.758217

**Authors:** Neelima Gupta, Aditya Verma, Prakhya Khandelwal, Tavpritesh Sethi

## Abstract

Machine learning models for molecular discovery often train on bioactivity data from heterogeneous assays, yet assay context is rarely explicitly represented. We investigate whether natural-language assay descriptions provide predictive information beyond molecular and protein representations. We develop ACL-DTI (Assay-aware Contrastive Learning-based Drug Target Interaction) and MAMMAL-AsAP, an assay-context extension of MAMMAL-DTI. Across five triplet-disjoint random splits, assay-aware models achieve competitive or improved performance over assay-agnostic baselines. Among the evaluated models, MAMMAL-AsAP achieves the highest (R^2^) of 0.7198 ± 0.0085. Masking experiments show that ACL-DTI is robust to incomplete assay descriptions, exhibiting the smallest reductions in R² among the evaluated models when masking the target name, assay technique, or both: 0.0228 ± 0.0064, 0.0406 ± 0.0113, and 0.0620 ± 0.0190, respectively. These results indicate that assay descriptions contain predictive information not fully captured by molecular structures and protein sequences, and reveal architecture-dependent differences in how this information is exploited.

---

Drug discovery frequently begins by determining whether a small molecule interacts with a target protein and, subsequently, estimating the strength of that interaction^1^. Traditionally, these tasks have relied on experimental assays that provide reliable mechanistic evidence but require substantial time, specialized resources, and financial investment^2,3^. As the number of available chemical compounds and biological targets continues to increase, the limitations of exclusively experimental screening have become increasingly evident. Consequently, computational approaches have emerged as efficient and scalable alternatives that enable large-scale exploration of drug-target relationships, prioritize candidate molecules, and support early-stage drug discovery while reducing reliance on exhaustive experimental testing. In this context, in-silico DTI prediction methods, particularly those based on deep learning (DL), have attracted considerable attention due to their potential to reduce development costs, accelerate drug discovery, and improve the success rate of candidate therapeutics^4^.

Bioactivity prediction underpins many applications in molecular machine learning, yet the bioactivity datasets used to train predictive models are inherently heterogeneous. Measurements for the same compound-target pair are often generated using different assay protocols, readouts, and experimental conditions, resulting in substantial assay-dependent variability. Historically, this challenge has been addressed during data curation by selecting a single representative measurement rather than preserving assay-specific observations^5^. Consequently, most molecular machine learning models remain effectively assay-agnostic, learning from pooled bioactivity measurements without explicitly modeling the experimental context in which they were generated. Recent work has emphasized that assay context is a major determinant of predictive performance and has advocated for assay-aware modeling strategies^6^.

Recent advances further demonstrate that natural-language assay descriptions encode biologically meaningful information. Incorporating assay descriptions alongside molecular representations improves molecular property prediction^7^, while *Assay2Mol: Large Language Model-based Drug Design Using BioAssay Context* leverages assay context for conditional molecular generation^8^. Despite these advances, existing approaches either focus on assay-aware data curation, couple assay descriptions only with molecular representations, or exploit assay context for generative tasks. A unified multimodal framework that explicitly models experimental context as a first-class modality alongside molecular structures and protein sequences for bioactivity prediction remains underexplored. Addressing this gap offers the potential to improve robustness, biological interpretability, and generalization across heterogeneous experimental settings.

To address this gap, we introduce *ACL-DTI – Assay-aware Contrastive Learning – Drug Target Interaction*, a contrastive learning-based multimodal framework that explicitly models experimental context by jointly learning representations of molecular structures, protein sequences, and natural-language assay descriptions for binding affinity prediction (Fig. 1). To demonstrate that assay-aware learning generalizes beyond a single architecture, we further extend IBM’s *MAMMAL* foundation model^9^ with assay representations, resulting in an assay-aware variant, *MAMMAL-AsAP* that incorporates experimental context as an additional modality. We systematically benchmark both assay-aware and assay-agnostic models spanning diverse architectural families to quantify the contribution of assay information across heterogeneous evaluation settings. Furthermore, through progressive masking of assay descriptions and comprehensive modality ablation studies, we investigate the robustness, interpretability, and relative importance of experimental context in multimodal representation learning. Together, our results demonstrate that assay context constitutes an independent and complementary source of predictive information, establishing experimental context as a first-class modality for molecular machine learning and providing a general framework for context-aware bioactivity prediction.

**Figure 1:**
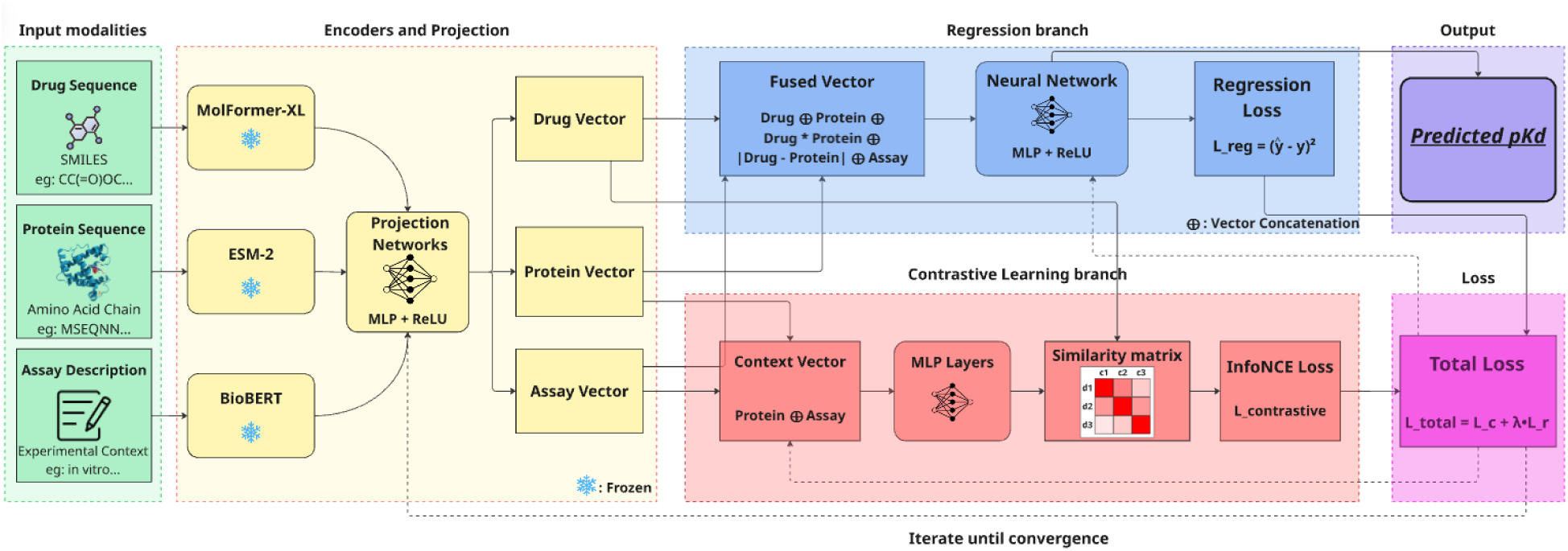
ACL-DTI framework for pK_d_ prediction with contrastive representation learning. The model integrates three input modalities: drug molecular structures represented as SMILES, protein sequences, and assay descriptions. Drug, protein, and assay inputs are encoded using MoLFormer-XL^10^, ESM-2^11^, and BioBERT^12^, respectively, with the pretrained encoders kept frozen and followed by modality-specific projection networks. The resulting drug, protein, and assay representations are used in two learning branches. In the regression branch, drug and protein representations are fused with the assay representation and passed to a multilayer perceptron to predict the drug pK_d_, with the regression loss defined as the squared prediction error. In parallel, the protein and assay representations are combined to construct a contextual representation, which is optimized using an InfoNCE-based contrastive loss^13^. The total training objective combines the regression and contrastive losses, and the model parameters are iteratively optimized until convergence.

**Figure 2:**
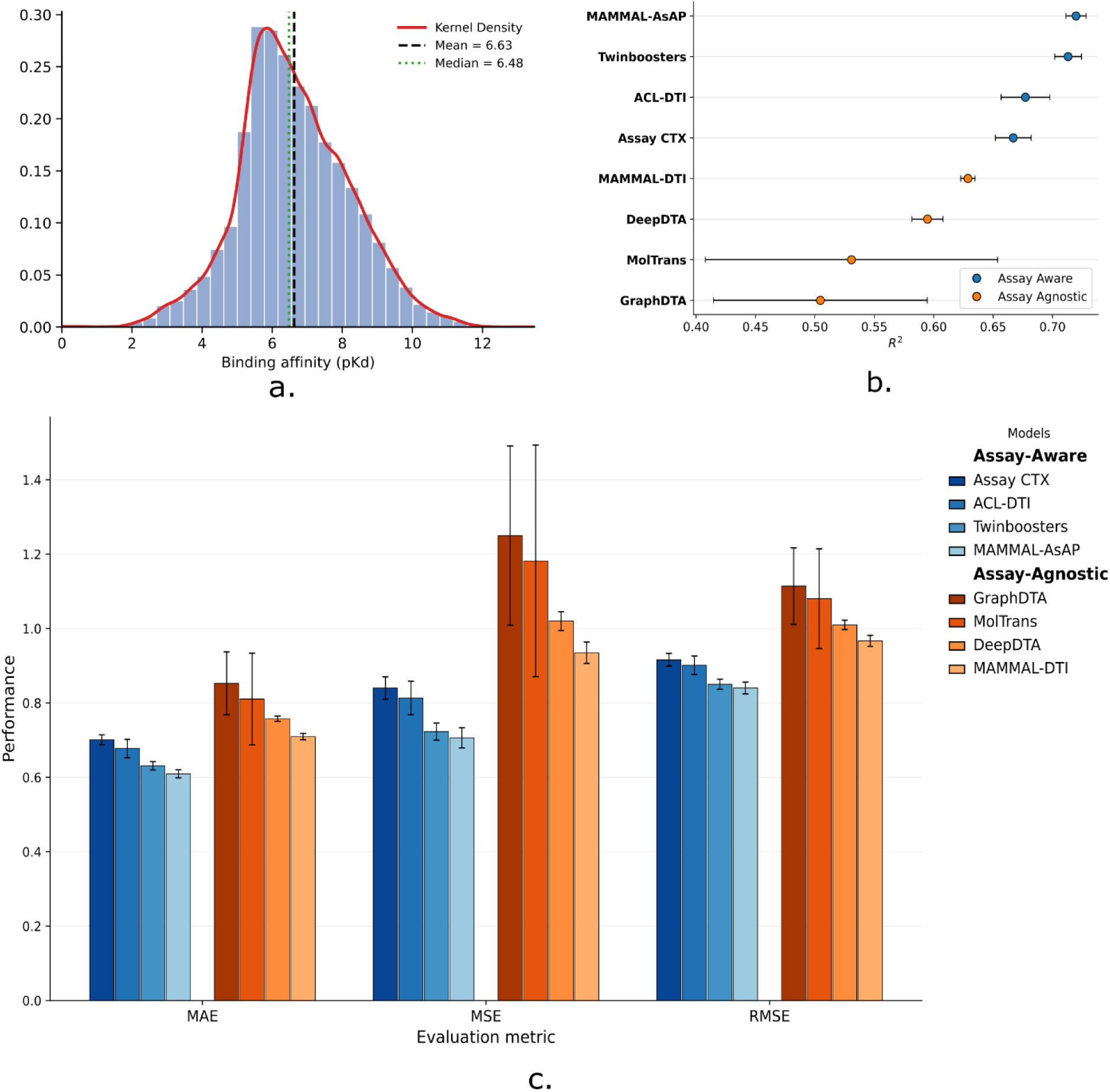
Assay-aware models consistently outperform assay-agnostic baselines. **a.** Kernel density estimate (KDE) of pK_d_ values across the complete dataset, illustrating the empirical distribution of the target variable. **b.** Cleveland dot plot comparing the mean R^2^ of assay-aware and assay-agnostic models across five independently generated triplet-disjoint data splits. Points denote the mean performance of each model, and horizontal error bars represent ±1 standard deviation across the five independent runs. Models are grouped by learning paradigm and ordered by decreasing mean R^2^ within each group, highlighting the consistent separation in predictive performance between assay-aware and assay-agnostic approaches. **c.** Mean MAE, MSE, and RMSE of assay-aware and assay-agnostic models across five independently generated triplet-disjoint data splits. Bars represent the mean performance across the five independent runs, with error bars denoting ±1 standard deviation. Lower values indicate better predictive performance. Assay-aware models show lower prediction errors across all three evaluation metrics than the assay-agnostic baselines.

## Results

### MAMMAL-AsAP improves drug-target binding-affinity prediction through assay-aware modeling

We first evaluated ACL-DTI and MAMMAL-AsAP for drug-target binding-affinity prediction using the curated ChEMBL dataset^14,15^. Performance on the held-out test set was assessed using the coefficient of determination (R²), mean absolute error (MAE), mean squared error (MSE) and root mean squared error (RMSE). Across the repeated evaluations, ACL-DTI achieved an R² of 0.6772 ± 0.0205, whereas MAMMAL-AsAP achieved an R² of 0.7198 ± 0.0085 (Fig. 2b). The corresponding MAE, MSE, and RMSE values are shown in Fig. 2c.

MAMMAL-AsAP therefore achieved a higher R² than ACL-DTI under the same evaluation setting. Because these models differ in multiple architectural components, the performance difference cannot be attributed to any individual component. We further compared MAMMAL-AsAP with MAMMAL-DTI (Fig. 2b,c), which provides a drug-target binding-affinity prediction setting without the assay-description modality. MAMMAL-AsAP achieved an R² of 0.7198 ± 0.0085, compared with 0.6288 ± 0.0060 for MAMMAL-DTI (Fig. 2b), together with lower error across the MAE, MSE and RMSE metrics (Fig. 2c). The higher R² and lower prediction errors observed for MAMMAL-AsAP are consistent with improved binding-affinity prediction when assay descriptions are incorporated. Because MAMMAL-AsAP and MAMMAL-DTI use the same underlying architecture and differ only in the incorporation of assay information, their performance difference provides a direct assessment of the contribution of assay descriptions to binding-affinity prediction.

### Assay information improves drug-target binding-affinity prediction

#### Assay-aware models consistently outperform assay-agnostic baselines

As shown in Fig. 2b, the assay-aware models achieved higher predictive performance than the assay-agnostic baselines across the evaluated architectures. All four assay-aware models achieved higher mean R² values than the assay-agnostic models, with MAMMAL-AsAP showing the highest predictive performance, followed by TwinBooster^7^. The proposed contrastive learning framework, ACL-DTI also outperformed the remaining assay-aware model, AssayCTX^6^. This performance difference was observed across all five independent data splits, with the corresponding variability reflected by the standard deviations of the reported metrics. Similar trends were observed for MAE, MSE and RMSE (Fig. 2c), indicating that the advantage of assay-aware models was not restricted to R² but was also reflected in prediction error across multiple evaluation metrics. The individual results for each split, including the bootstrap-derived estimates, are provided in the **Supplementary Information** (Supplementary Tables 2 and 3). The masking and ablation analyses presented in the following sections further examine the contribution of assay-derived information to these performance differences.

#### Masking assay information reveals model-specific dependence on assay context

The masking experiments revealed distinct, architecture-dependent sensitivities to assay-description information **(Fig. 3a**; **Table 1)**. AssayCTX^6^ showed the strongest dependence on the unmasked assay description, with target-name masking causing a pronounced loss of predictive performance; masking the assay technique alone produced a smaller effect, whereas simultaneous masking did not lead to a further appreciable reduction beyond target-name masking. Twinbooster^7^ exhibited a similar preferential sensitivity to target-name masking, although its response to combined masking was more pronounced. In contrast, ACL-DTI and MAMMAL-AsAP were comparatively stable under either individual masking perturbation. For these models, simultaneous masking produced a clearer reduction in performance than either individual perturbation, particularly for MAMMAL-AsAP**(Fig. 3a)**. Thus, the effect of removing assay-description information depended on both the model architecture and the specific semantic component removed, with no uniform response to the combined loss of target and assay information. Detailed results for all evaluation metrics across all random seeds are provided in **Supplementary Tables (5-10)**.

**Figure 3:**
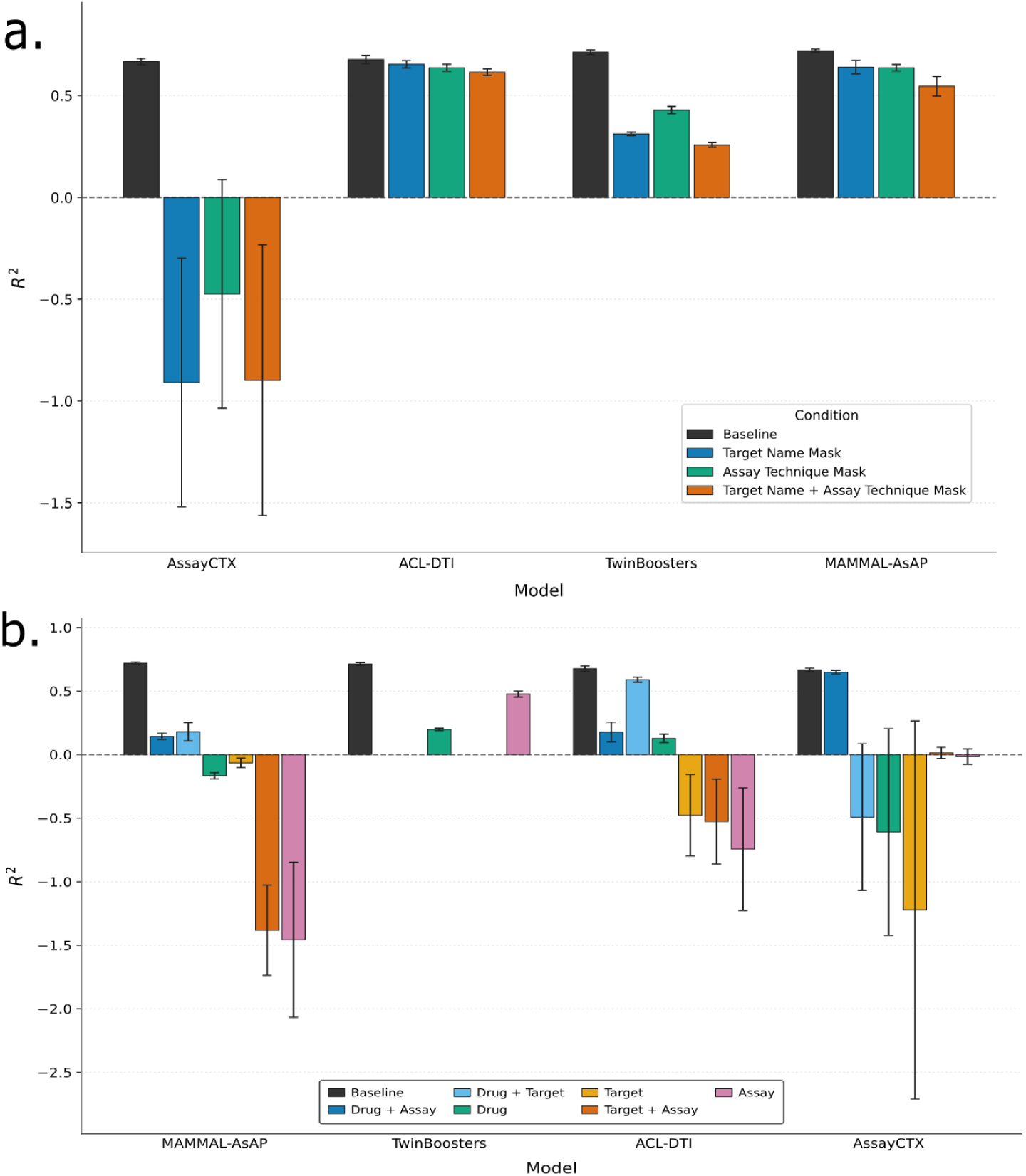
**a. Comparison of predictive performance (R^2^) across different evaluated architectures for all the masking conditions**. Bars represent the mean R^2^, and error bars indicate the corresponding ±1 standard deviation across the five independent runs. **b. Comparison of predictive performance (R^2^) across different evaluated architectures using different input modality combinations.** Bars represent the mean R^2^, and error bars indicate the corresponding ±1 standard deviation across the five independent runs.

These patterns suggest that the evaluated architectures differ in the extent to which their predictions depend on information encoded in assay descriptions. The pronounced sensitivity of AssayCTX^6^ and Twinbooster^7^ to target-name masking is consistent with greater dependence on target-related information, whereas the relative stability of ACL-DTI and MAMMAL-AsAP indicates that their predictive performance is less affected by these individual perturbations. However, the masking experiment does not distinguish whether the observed differences arise from differential use of target or assay information, differences in representation learning, or other architectural properties. In particular, the response to simultaneous masking does not follow a uniform additive pattern across models, motivating further controlled ablation or representation-level analyses.

#### Modality ablation reveals the contribution of individual input modalities

We next evaluated the contribution of individual modalities through systematic ablation of drug, target, and assay-description inputs. Removing assay descriptions reduced (*R*^2^) for all four models, although the magnitude of the reduction varied substantially. For AssayCTX^6^, (*R^2^*) decreased from (0.667 ± 0.015) to (–0.492 ± 0.577), whereas the corresponding reductions were smaller for ACL-DTI, from (0.6772 ± 0.0205) to (0.590 ± 0.020), and for MAMMAL-AsAP, from (0.7198 ± 0.0085) to (0.180 ± 0.072). In the case of Twinbooster^7^, removing the assay description drops performance from (0.7131 ± 0.0112) to (0.199 ± 0.010). Thus, assay descriptions provide predictive information in all four models, but their contribution depends on the model architecture and the information retained in the other modalities. In particular, the relatively small reduction in ACL-DTI indicates that much of its predictive performance can be retained without assay descriptions, whereas the larger reductions for MAMMAL-AsAP and AssayCTX^6^ indicate greater sensitivity to their removal. These results support a contribution of assay information to binding-affinity prediction without implying that it is uniformly the dominant modality.

The ablation patterns further reveal differences in modality dependence between models. For ACL-DTI and MAMMAL-AsAP, which learn joint representations from drug, target, and assay descriptions, removing the drug modality produced the largest reduction in (*R2*), followed by removing the target and assay-description modalities. Thus, for these models, the observed sensitivity to modality removal follows the order drug > target > assay description. AssayCTX^6^ showed a different pattern, with the largest reduction after removing assay descriptions, followed by drug and target. This difference suggests that modality contributions are coupled to the representation-learning strategy rather than being uniform across models. Twinbooster^7^ uses only drug and assay inputs and therefore permits comparison of only the corresponding single-modality conditions.

The single-modality experiments provide a complementary perspective on the role of each modality. For ACL-DTI and MAMMAL-AsAP, assay descriptions produced the lowest predictive performance when used in isolation, indicating that their contribution is not primarily reflected in standalone prediction. In contrast, AssayCTX^6^ achieved its highest single-modality performance using assay descriptions, whereas target-only input yielded the lowest performance among the three modalities. Together with the multimodal ablations, these results indicate that standalone predictive utility and incremental contribution to a multimodal representation are not equivalent properties.

Overall, these results highlight that the predictive value of a modality depends not only on its standalone information content but also on how it interacts with the other modalities within each model’s representation-learning framework.

## Discussion

Our results show that incorporating assay information consistently improves drug-target binding affinity prediction relative to corresponding assay-agnostic configurations, with the same trend observed across both neural architectures and the comparatively simple LGBM model. This suggests that assay descriptions provide predictive information beyond compound and target identity and that their utility is not restricted to complex multimodal architectures. Our first contribution, MAMMAL-AsAP, an extension of the MAMMAL architecture^9^ with assay description, achieved the highest overall performance. Twinbooster^7^ ranked second overall despite not incorporating target information. When considering models that use all three modalities, drug, target, and assay, our second contribution, the contrastive learning-based ACL-DTI, achieved the second-highest performance, ranking third overall. Although MAMMAL-AsAP combines modular tokenization and prompt-based transformer modeling, the present comparisons do not isolate the contribution of these individual components; therefore, its performance advantage should be interpreted as a property of the evaluated system rather than attributed to a specific architectural choice.

The masking and modality-ablation experiments further demonstrated that models differ in their dependence on assay and target information. The relatively weak target-only performance may reflect differences in protein representation, such as the use of Z-scale descriptors^16,17^ in AssayCTX^6^ compared with pretrained representations such as ESM-2^11^ in ACL-DTI, although protein representation and architecture were not independently controlled; therefore, this interpretation remains tentative.

Across models, drug information emerged as an important predictive modality, whereas the relative contributions of target and assay information varied by modeling framework. The stronger performance of complete multimodal configurations compared with many ablated configurations supports the complementary value of molecular, target, and assay information. However, the evaluated models differ in architecture, input representations, training objectives, and fusion mechanisms, preventing attribution of these effects to any individual component of the model. Likewise, masking evaluates sensitivity to missing information rather than the robustness of learned representations. The comparison should also be interpreted within the available modality space: for example, Twinbooster^7^ does not include a target modality and therefore cannot be directly compared with three-modality architectures in target-specific ablations.

Overall, these findings support assay-aware modeling as a useful direction for drug-target affinity prediction and indicate that assay context can contribute predictive information across substantially different modeling approaches. At the same time, the study is based on ChEMBL-derived measurements^14,15^, whose assay descriptions and experimental conditions are heterogeneous, and the extent to which these findings generalize to independent datasets remains to be established. Controlled experiments that independently vary modality representations and fusion mechanisms, together with external or prospective experimental validation, will be important for determining whether the observed gains reflect transferable assay-context information rather than dataset-specific associations.

More broadly, the results support the explicit integration of experimental context into multimodal molecular machine-learning models. Future work should assess the generalization of assay-aware models across unseen assay contexts and under incomplete or distribution-shifted experimental information. Improving the standardization and quality of assay metadata, together with developing models that can jointly represent molecular, protein, and experimental context, may improve the reliability and generalizability of computational affinity prediction and support more effective prioritization of compounds for experimental validation.

## Methods

### Dataset Curation and Splitting

Bioactivity data were retrieved from ChEMBL release 35 using the ChEMBL Web API^14,15^. To reduce ambiguity associated with censored measurements, only records reporting dissociation constants (K_d_) with an exact relation (“=”) were retained, while entries annotated with inequality operators (e.g., “<” or “>”) were excluded. The dataset was further restricted to measurements reported in nM for *Homo sapiens* single-protein targets. Molecular structures represented as Simplified Molecular Input Line Entry System (SMILES) strings were standardized by converting them to canonical SMILES using RDKit^18^. Binding affinities were transformed from (K_d_) to (pK_d_) according to:

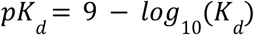

where (K_d_) is expressed in nM. To improve computational efficiency, maximum input lengths were defined for each modality. Drug representations were limited to 250 SMILES characters, protein sequences to 1,250 amino acids, and assay descriptions to 300 characters. Samples exceeding these limits were excluded from the dataset. The distribution of (pK_d_) values is illustrated in **Fig. 2a**.

The curated dataset was partitioned into training, validation, and test sets using an 80:10:10 ratio. A triplet-disjoint splitting strategy was adopted, whereby each unique drug-target-assay combination was assigned exclusively to one partition. This ensured that identical interaction records did not appear across multiple data splits.

### ACL-DTI Model Architecture

To effectively capture the complex interplay among compound structure, protein sequence, and experimental context, we developed a tri-modal deep learning architecture that leverages three independent, pre-trained foundation models. Specifically, compound SMILES strings were encoded using MoLFormer-XL^10^ to extract structural topology, target amino acid sequences were processed with ESM-2^11^ to capture evolutionary and functional characteristics, and experimental assay descriptions were encoded using BioBERT^12^ to interpret complex contextual protocols. The raw representations extracted from these modality-specific encoders were subsequently passed through independent Multi-Layer Perceptron (MLP) projection heads, systematically mapping them into a shared 512-dimensional continuous latent space.

To optimize the architecture, we implemented a composite loss function (*L_total_* = *L_contrastive_* + λ *L_regression_*) that simultaneously aligns latent manifolds and predicts continuous binding affinities. For robust contrastive learning, a specialized Context Encoder fused the un-normalized protein and assay projections into a cohesive “assay-aware” context vector; this vector and the projected drug representation were *L_2_* normalized and geometrically aligned via a Symmetric InfoNCE loss^13^, systematically pulling interacting triplets together in the latent space while repelling non-interacting samples. Concurrently, the final scalar pKd prediction was generated by concatenating the normalized drug vector, normalized protein vector, their explicit interaction synergy (Hadamard product), their latent space distance (absolute difference), and the un-normalized raw assay representation into a comprehensive 2560-dimensional joint embedding, which was then processed through a multi-layer Fusion Regressor network optimized via Mean Squared Error (MSE) loss (Fig. 1).

### MAMMAL-AsAP Model Architecture

MAMMAL-AsAP is built upon the pretrained MAMMAL foundation model^9^ and adapts its multimodal architecture to downstream drug-target binding-affinity prediction. MAMMAL employs a transformer-based architecture with a Modular Tokenizer and a prompt-based input formulation, enabling heterogeneous molecular and biological entities to be represented within a unified sequence framework^9^. The pretrained model was used as the initialization for MAMMAL-AsAP, and its parameters were subsequently fine-tuned on the curated binding-affinity dataset.

For binding-affinity prediction, MAMMAL-AsAP jointly incorporates three inputs: the drug, the target protein, and the corresponding assay description. Drug and target sequences were represented using MAMMAL’s molecular and protein tokenization components^9^, respectively, while assay descriptions were incorporated as additional textual input via the Modular Tokenizer and prompt-based input formulation. These inputs were jointly fed into the MAMMAL encoder, enabling the model to learn representations of the drug-target interaction in the context of the experimental assay.

The fine-tuned model predicts binding affinity as a continuous-valued output using MAMMAL’s regression capability. During downstream training, the MAMMAL parameters were updated jointly using the three input modalities and the corresponding experimentally measured binding-affinity values. Thus, MAMMAL-AsAP retains the pretrained representation and multimodal modeling framework of MAMMAL^9^ while adapting the model to incorporate assay-specific contextual information for binding-affinity prediction.

#### Benchmarking Strategy

To evaluate the contribution of assay information to bioactivity prediction, we benchmarked four assay-aware models and four assay-agnostic baselines. The selected methods encompass diverse modeling paradigms, including multimodal deep learning, transformer-based architectures, graph neural networks, proteochemometric approaches, and gradient boosting methods, providing a broad representation of current strategies for drug-target affinity prediction.

#### Assay-Aware Models

In addition to the two models described above, we implemented and evaluated two additional assay-aware models as benchmarks for binding-affinity prediction, both of which incorporate assay information as input.

#### AssayCTX

AssayCTX is a proteochemometric framework that incorporates ligand descriptors, protein descriptors, and assay information for bioactivity prediction^6^. Following the original implementation, assay descriptions were represented using the BioBERT-derived embeddings reported by the authors to provide assay-specific contextual information. The resulting representations were used within the original LightGBM-based prediction framework^6^.

#### TwinBooster

TwinBooster integrates molecular representations and assay embeddings using a multimodal representation-learning framework^7^. Molecular features were generated from extended-connectivity fingerprints (ECFPs)^19^, whereas assay descriptions were encoded using the pretrained DeBERTa model^20^ released with the original implementation^7^. The learned joint representations were subsequently used to train a LightGBM regressor for continuous (pK_d) prediction following the original training protocol.

#### Assay-Agnostic Models

To provide a direct comparison with methods that do not utilize assay information, four assay-agnostic models were evaluated using the same training, validation, and test partitions.

### MAMMAL-DTI

MAMMAL-DTI is the assay-agnostic variant of the MAMMAL framework that predicts binding affinity using SMILES sequence and protein sequences without incorporating assay descriptions^9^. The pretrained MAMMAL model was fine-tuned for binding-affinity prediction using a ChEMBL-derived dataset curated for this study. Although the MAMMAL authors provide a drug-target interaction (DTI) model fine-tuned on BindingDB^21^, we did not use it in our downstream experiments. Instead, we initialized MAMMAL-DTI from the pretrained MAMMAL checkpoint and fine-tuned it on the ChEMBL dataset^14,15^ using drug, target, and assay-description information as model inputs.

#### DeepDTA

DeepDTA is a sequence-based deep learning model that predicts binding affinity from drug SMILES and protein sequences using convolutional neural networks^22^. The original preprocessing pipeline and model architecture were adopted for benchmarking.

#### MolTrans

MolTrans employs transformer-based encoders to learn contextual representations of molecular and protein substructures before modeling cross-modal interactions for affinity prediction^23^. The original tokenization strategy, model architecture, and preprocessing pipeline were used.

#### GraphDTA

GraphDTA represents compounds as molecular graphs and proteins as amino acid sequences to predict binding affinity using graph neural networks^24^. Drug molecules were converted from SMILES strings to molecular graphs as in the original implementation, while protein sequences were processed with the corresponding sequence encoder.

All benchmark models were trained and evaluated using the identical data partitions and evaluation protocol described in the section ‘Data Curation and Splitting’ to enable a consistent comparison between assay-aware and assay-agnostic approaches.

#### Multi-Seed Evaluation

To account for variability arising from data partitioning, all models were evaluated across five independently generated triplet-disjoint data splits.

For each data split, regression performance metrics were estimated by bootstrap resampling of the test-set predictions with 1,000 iterations. The bootstrap distributions were used to compute the mean performance metric and the corresponding 95% confidence interval for each seed; all results are reported in the **Supplementary Results**.

Overall model performance was then summarized by averaging the point estimates obtained from the five independent data splits. To quantify variability arising from different data partitions, the standard deviation of the performance metrics across the five seeds was also reported. Thus, bootstrap confidence intervals characterize the uncertainty associated with each test set, whereas the variability across seeds reflects the robustness of model performance under different train-validation-test partitions.

### Evaluation Strategy

#### Progressive Assay Description Masking

To quantify the contribution of assay-derived contextual information, assay-aware models were evaluated under progressively masked assay descriptions. Three masking strategies were considered: (i) masking target names explicitly mentioned in the assay description, (ii) masking assay-specific experimental descriptors, including protocols and experimental techniques, and (iii) masking both target names and assay-specific descriptors.

Target names and Assay techniques were identified using the GLiNER Biomedical named-entity recognition model^25^. Entity mentions classified as target names, including recognized synonyms, were masked while preserving the remaining assay description. No additional manual preprocessing or tokenization was performed following masking; the modified descriptions were processed using each model’s standard input pipeline. All masking experiments were conducted exclusively on the test partitions of the five triplet-disjoint data splits to evaluate the dependence of assay-aware models on different categories of assay-derived information.

#### Ablation Study

An ablation study was conducted to quantify the contribution of each input modality to binding affinity prediction. Six input configurations were evaluated: (i) drug and protein sequence, (ii) drug and assay description, (iii) protein sequence and assay description, (iv) drug only, (v) protein sequence only, and (vi) assay description only.

For each configuration, the model architecture, training procedure, and hyperparameters were kept unchanged. The excluded modality was removed by replacing the corresponding input field with an empty value, thereby preserving the original model architecture while eliminating information from the omitted modality.

## Data Availability

The data used in this study were obtained from the publicly available ChEMBL database and accessed using the ChEMBL web source client^14,15^. The data can be accessed directly from the ChEMBL database at https://www.ebi.ac.uk/chembl/.

## Code Availability

The ACL-DTI and MAMMAL-AsAP frameworks, along with all scripts required to reproduce the analyses, are available at https://github.com/tavlab-iiitd/AsAP. The code is released under the MIT License.

## Author Contributions

NG– (Conceptualization [equal], Data curation [equal], Software [equal], Visualization [equal], Writing: original draft [equal], Writing: review & editing [equal]); AV– (Software [equal], Writing: original draft [equal], Writing: review & editing [equal]); PK– (Software [supporting]); TS– (Conceptualization [equal], Methodology [equal], Supervision [lead], review & editing [equal])

All authors read and approved the final version of the manuscript and agree to be accountable for all aspects of the work.

## Acknowledgments

The authors acknowledge financial support from the Department of Biotechnology (DBT), Ministry of Science and Technology, Government of India, under project BT/PR40195/BTIS/137/58/2023 provided to ACBR and IIIT-Delhi. We thank the Department of Computational Biology, IIIT-Delhi, for providing computational and infrastructural support. We also acknowledge Dr. Madhu Chopra (ACBR) for her support during the course of this work.

## Tables

**Table 1:** Predictive performance of multimodal models under assay-description masking. Model performance was evaluated under three masking conditions: target-name masking, assay-technique masking, and combined target-plus-assay masking. Changes in predictive performance are reported as ΔR² relative to the unmasked condition, with more negative values indicating larger reductions in R² following masking. Values are reported as mean ± s.d.

| Model | Target name Masked ($\Delta R^2$) | Assay Masked ($\Delta R^2$) | Target+Assay Masked ($\Delta R^2$) |
| --- | --- | --- | --- |
| MAMMAL-AsAP | $-0.0800 \pm 0.0345$ | $-0.0830 \pm 0.0185$ | $-0.1735 \pm 0.0522$ |
| ACL-DTI | $-0.0228 \pm 0.0064$ | $-0.0406 \pm 0.0113$ | $-0.0620 \pm 0.0190$ |
| AssayCTX | $-1.5760 \pm 0.6161$ | $-1.1401 \pm 0.5633$ | $-1.5651 \pm 0.6698$ |
| Twinbooster | $-0.4006 \pm 0.0101$ | $-0.2844 \pm 0.0162$ | $-0.4556 \pm 0.0143$ |

**Table 2:** Ablation analysis of input modalities across all evaluated architectures. The table reports the change in predictive performance (ΔR²) relative to the corresponding reference model after removing individual or combined input modalities. Each ablation configuration retains only the listed input modalities. Larger reductions in R² indicate a greater contribution of the omitted modality to overall predictive performance. Values are reported as the mean ΔR² with ±1 standard deviation across the five independent runs.

| Model | Input configuration |  |  |  |  |  |
| --- | --- | --- | --- | --- | --- | --- |
| | Target + Assay<br>($\Delta R^2$) | Drug + Target<br>($\Delta R^2$) | Drug+Assay<br>($\Delta R^2$) | Assay<br>($\Delta R^2$) | Drug<br>($\Delta R^2$) | Target<br>($\Delta R^2$) |
| ACL-DTI | $-1.2040 \pm 0.3361$ | $-0.0875 \pm 0.0061$ | $-0.4993 \pm 0.0948$ | $-1.4214 \pm 0.4983$ | $-0.5497 \pm 0.0474$ | $-1.1546 \pm 0.3244$ |
| MAMMAL-AsAP | $-2.1017 \pm 0.3545$ | $-0.5401 \pm 0.0702$ | $-0.5761 \pm 0.0258$ | $-2.1770 \pm 0.6106$ | $-0.8863 \pm 0.0315$ | $-0.7836 \pm 0.0318$ |
| AssayCTX | $-0.6525 \pm 0.0440$ | $-1.1591 \pm 0.5758$ | $-0.0180 \pm 0.0019$ | $-0.6828 \pm 0.0689$ | $-1.2755 \pm 0.8099$ | $-1.8886 \pm 1.4825$ |
| Twinbooster | - | - | $0.0000 \pm 0.0000$ | $-0.2360 \pm 0.0249$ | $-0.5141 \pm 0.0152$ | - |

## Supplementary

### Supplementary Methods

#### 1. ACL-DTI Methodology

##### 1.1 Overview

ACL-DTI (Assay-aware Contrastive Learning for Drug Target Interaction) is a three-modality, dual-branch deep learning pipeline. Three separate pretrained encoders convert drug SMILES, protein sequences, and assay descriptions into dense vector representations. These representations are then processed by two branches that run in parallel throughout training: a contrastive learning branch that aligns the drug embedding with an assay-fused protein context in a shared latent space, and a regression branch that directly predicts pKd from the concatenation of all three modality embeddings together with drug-protein interaction and dissimilarity features. Both branches share the same projection layers and are trained jointly via a combined loss. The complete pipeline is illustrated in Fig 1.

**Figure 1:**
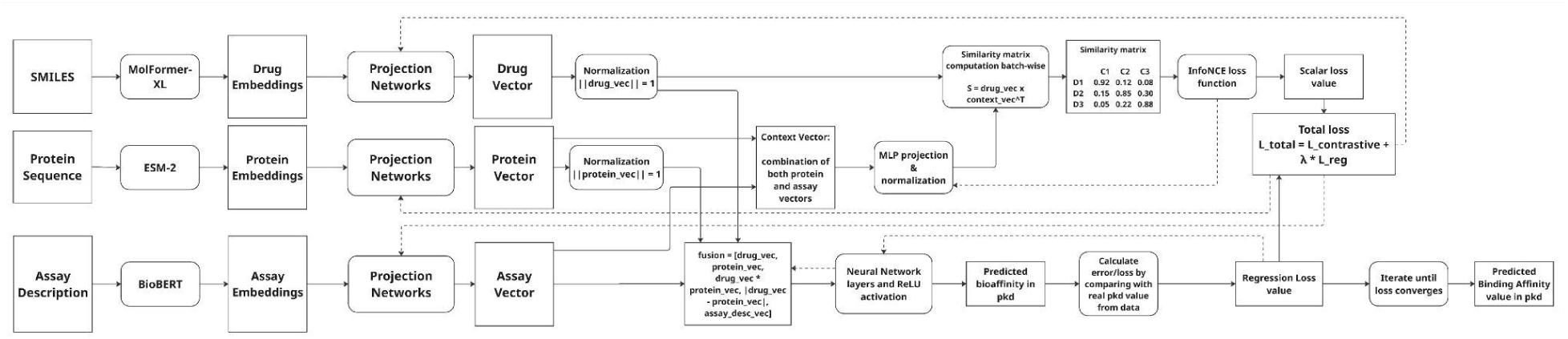
ACL-DTI Architecture. Drug, protein, and assay embeddings are each projected to 512 dimensions. Protein and assay projections are fused into an assay-conditioned context vector, which forms one pole of the symmetric InfoNCE contrastive loss1. In parallel, all three projected embeddings are concatenated – along with the element-wise product and absolute difference of the drug and protein embeddings – and passed through a regression MLP to predict pKd. Both branches are trained simultaneously via a combined loss.

##### 1.2 Multi-Modal Encoding

The first stage converts each of the three raw inputs into a fixed-length embedding vector using domain-specific pretrained language models.

###### 1.2.1 Drug Encoder: MoLFormer-XL

MoLFormer-XL^2^ encodes the drug’s SMILES string. Its linear-attention transformer architecture enables efficient processing of long chemical sequences, and its pretraining on 1.1 billion SMILES strings from

ZINC and PubChem endows it with broad coverage of chemical space. Mean pooling over token embeddings yields a 768-dimensional molecular representation. The encoder weights are kept frozen throughout training. SMILES strings are capped at 512 tokens; longer molecules are truncated.

###### 1.2.2 Protein Encoder: ESM-2

ESM-2^3^ encodes the protein amino acid sequence. The 35M-parameter variant (esm2_t12_35M_UR50D) is used; this size was chosen to fit within available GPU memory constraints; larger ESM-2 variants may yield richer residue-level representations but were not explored here. This produces 480-dimensional residue-level embeddings. Mean pooling over residue embeddings yields a compact protein-level representation that captures evolutionary and structural information encoded in large-scale sequence pretraining. The encoder is frozen. Sequences are capped at 1022 residues; longer proteins are truncated.

###### 1.2.3 Assay Encoder: BioBERT

BioBERT^4^ encodes the natural-language assay description. Assay descriptions (e.g., “Radioligand binding assay measuring inhibition of [3H]-citalopram binding to human SERT at 4nM”) convey the experimental modality, detection method, and measurement conditions. BioBERT’s^4^ pretraining on PubMed and PubMed Central^5^ text makes it sensitive to the biomedical terminology in these descriptions. The [CLS] token embedding (768-dimensional) is used as the assay representation. The encoder is frozen. Assay descriptions are capped at 300 tokens; longer descriptions are truncated.

###### 1.2.4 Projection heads

Because each encoder produces embeddings of different dimensionality and from different representational spaces, all three representations are independently projected to a shared latent dimension of 512 via a two-layer MLP:

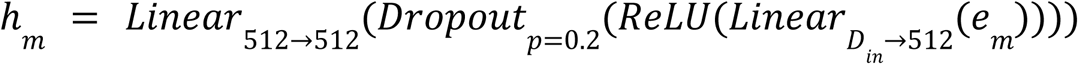

Where *e_m_* is the frozen-encoder embedding of modality *m* ∈ {drug, protein, assay}, *Din_d_* = 768, *Din_p_* = 480, *Din_a_* = 768 and a dropout of *p* = 0. 2 is applied within each head. Drug and protein projections are L2-normalised, whereas the assay representation is retained without L2-normalization:

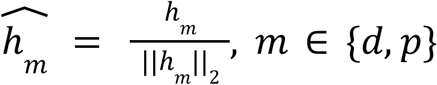

##### 1.3 Assay-Fused Context Construction

A core design requirement of this pipeline is that the assay information must enter the contrastive objective so that the shared latent space is conditioned on experimental context. The InfoNCE loss^1^ accepts exactly two inputs, so the protein and assay projections are fused into a single context vector ĉ before the contrastive branch. Specifically, the two projected vectors are concatenated into a 1024-dimensional vector, which is projected back to 512 dimensions and L2-normalised:

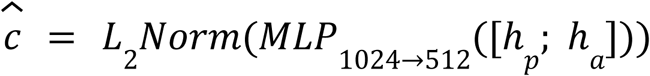

This context vector encodes both what the protein is (from ESM-2^3^) and how it was measured (from BioBERT^4^), making the contrastive alignment sensitive to assay conditions.

##### 1.4 Contrastive Learning Branch

The contrastive branch aligns drug representations with their corresponding assay-conditioned protein contexts. For a batch of B samples, the drug vectors {ĥ*_d_*_,*i*_} and context vectors {ĉ*_i_*} form B positive pairs.

All other within-batch pairings are treated as implicit negatives. The pairwise similarity matrix is:

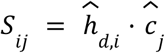

The symmetric InfoNCE loss^1^ is then:

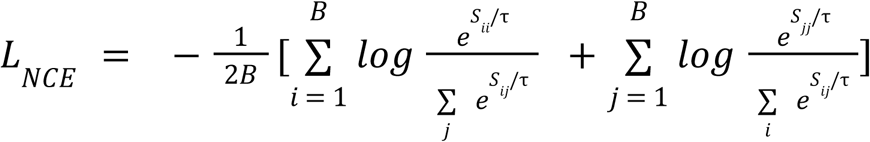

with temperature τ = 0.1. By operating symmetrically in both drug-to-context and context-to-drug directions, the loss enforces mutual alignment. The temperature τ controls the sharpness of the distribution; smaller values sharpen the contrast between positives and negatives.

##### 1.5 Regression Branch

The regression branch predicts the pKd scalar directly from a rich feature vector assembled from all three modalities. In addition to the three projected embeddings, the element-wise product of drug and protein vectors *ĥ_d_* ⊙ *ĥ_p_* (interaction features) and their element-wise absolute difference |*ĥ_d_* − *ĥ_p_* | (dissimilarity features) are appended, yielding a 2560-dimensional input to the MLP:

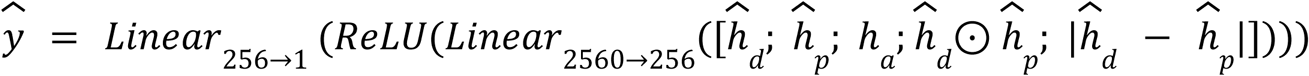

The MSE regression loss is:

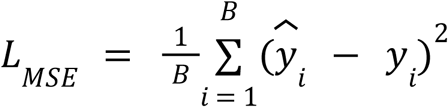

Where *ŷ_i_* and *y_i_* denote the predicted and experimental pKd of sample *i*, respectively.

##### 1.6 Joint Training and Combined Loss

Both branches run in parallel: neither is pretrained separately, and neither is frozen while the other trains. At every training step, forward passes through both branches are computed simultaneously, producing *L_NCE_* and *L_MSE_* simultaneously. The combined loss:

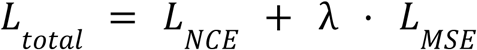

with λ = 0.1 (The weighting coefficient was selected empirically to balance the relative magnitudes of the two loss terms) is then back-propagated in a single pass, updating the projection heads, the context fusion layer, and the regression MLP jointly. This parallel design ensures that the contrastive alignment and the regression objective mutually inform each other throughout training: the contrastive branch shapes the latent geometry to reflect biochemical interaction, while the regression branch steers that geometry toward accurate affinity prediction. The choice to freeze all pretrained encoders reflects both a practical constraint (available GPU memory) and a principled one: the pretrained representations already encode rich chemical, biological, and biomedical knowledge, and fine-tuning with a relatively small dataset risks catastrophic forgetting of that knowledge.

#### 2. Benchmarked Architectures and Adaptations

**Table 1:**
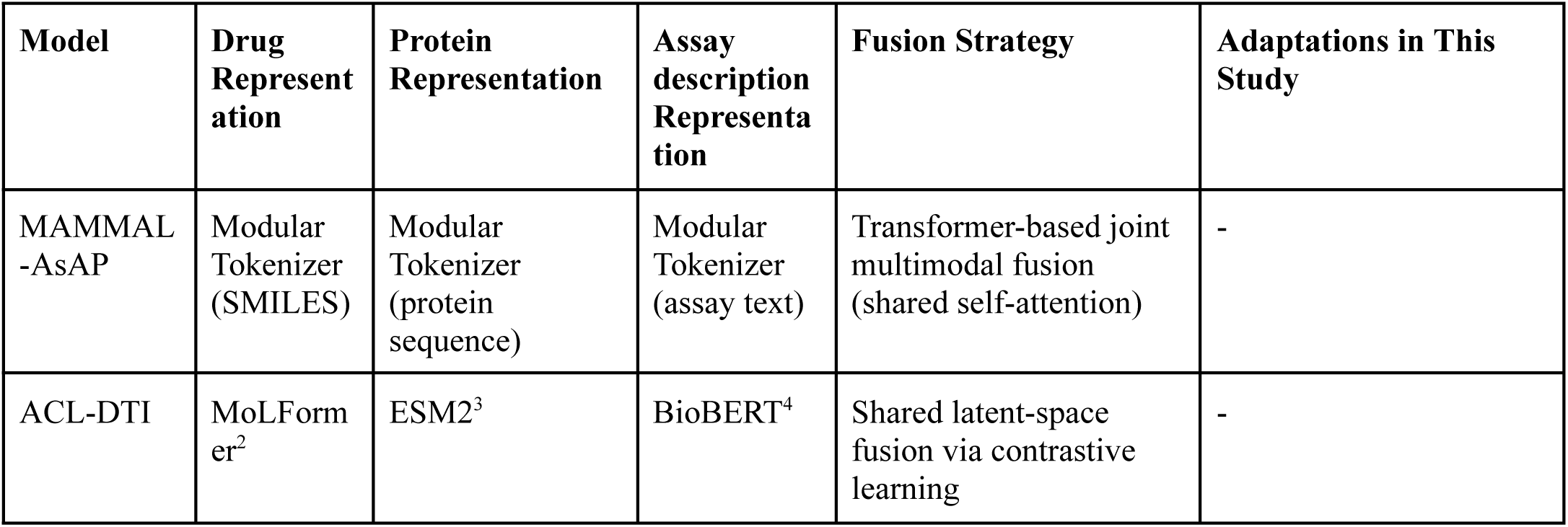
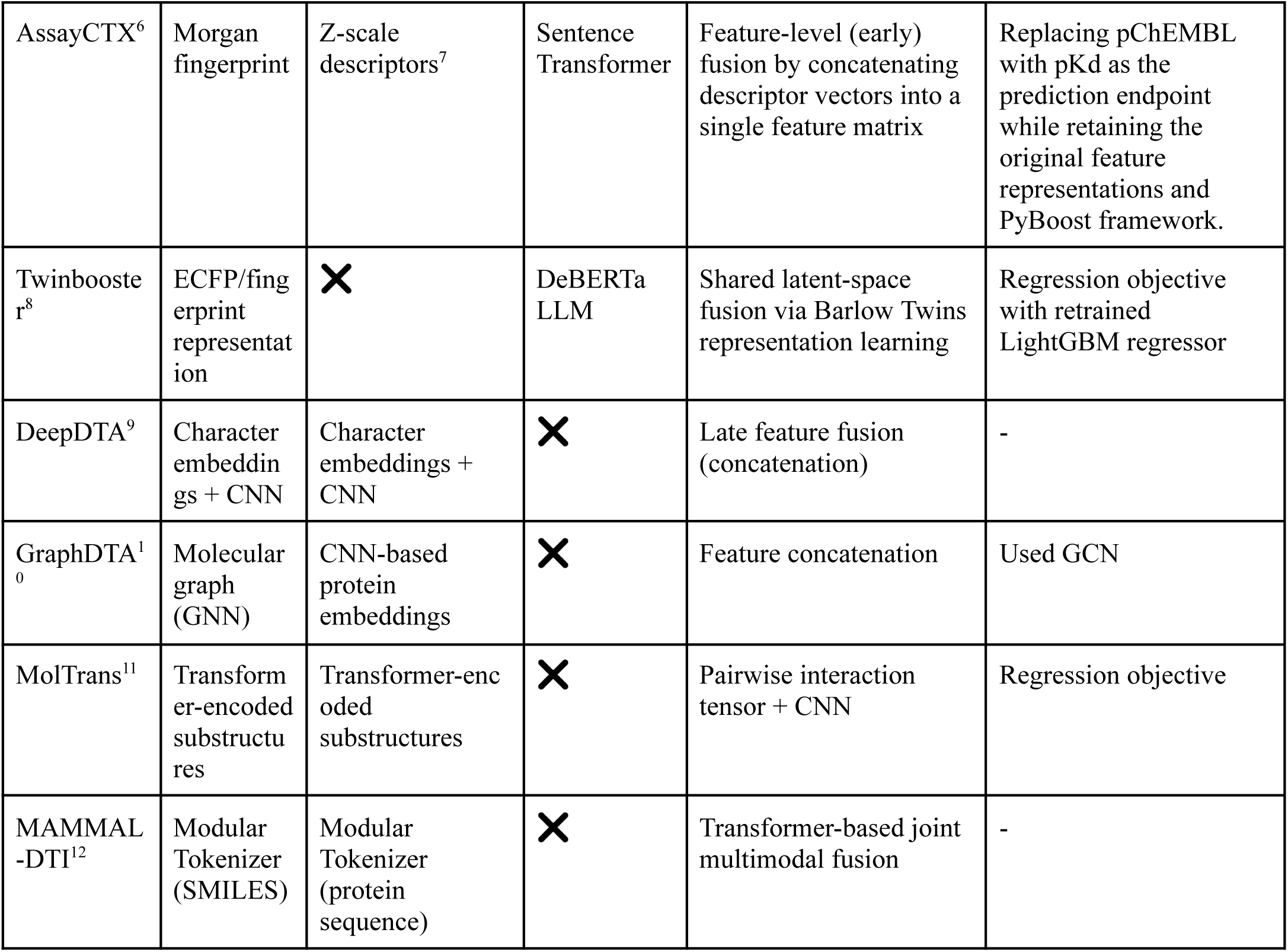
Architectural comparison of the benchmarked models, highlighting the representations used for each input modality (drug, protein, and assay, where applicable) and their corresponding multimodal fusion strategies.

| Model | Drug Representation | Protein Representation | Assay description Representation | Fusion Strategy | Adaptations in This Study |
| --- | --- | --- | --- | --- | --- |
| MAMMAL-AsAP | Modular Tokenizer (SMILES) | Modular Tokenizer (protein sequence) | Modular Tokenizer (assay text) | Transformer-based joint multimodal fusion (shared self-attention) | - |
| ACL-DTI | MolFormer<sup>2</sup> | ESM2<sup>3</sup> | BioBERT<sup>4</sup> | Shared latent-space fusion via contrastive learning | - |
| AssayCTX<sup>6</sup> | Morgan fingerprint | Z-scale descriptors<sup>7</sup> | Sentence Transformer | Feature-level (early) fusion by concatenating descriptor vectors into a single feature matrix | Replacing pChEMBL with pKd as the prediction endpoint while retaining the original feature representations and PyBoost framework. |
| Twinbooster<sup>8</sup> | ECFP/fingerprint representation | × | DeBERTa LLM | Shared latent-space fusion via Barlow Twins representation learning | Regression objective with retrained LightGBM regressor |
| DeepDTA<sup>9</sup> | Character embeddings + CNN | Character embeddings + CNN | × | Late feature fusion (concatenation) | - |
| GraphDTA<sup>10</sup> | Molecular graph (GNN) | CNN-based protein embeddings | × | Feature concatenation | Used GCN |
| MolTrans<sup>11</sup> | Transformer-encoded substructures | Transformer-encoded substructures | × | Pairwise interaction tensor + CNN | Regression objective |
| MAMMAL-DTI<sup>12</sup> | Modular Tokenizer (SMILES) | Modular Tokenizer (protein sequence) | × | Transformer-based joint multimodal fusion | - |

## Supplementary Results

**Figure 2:**
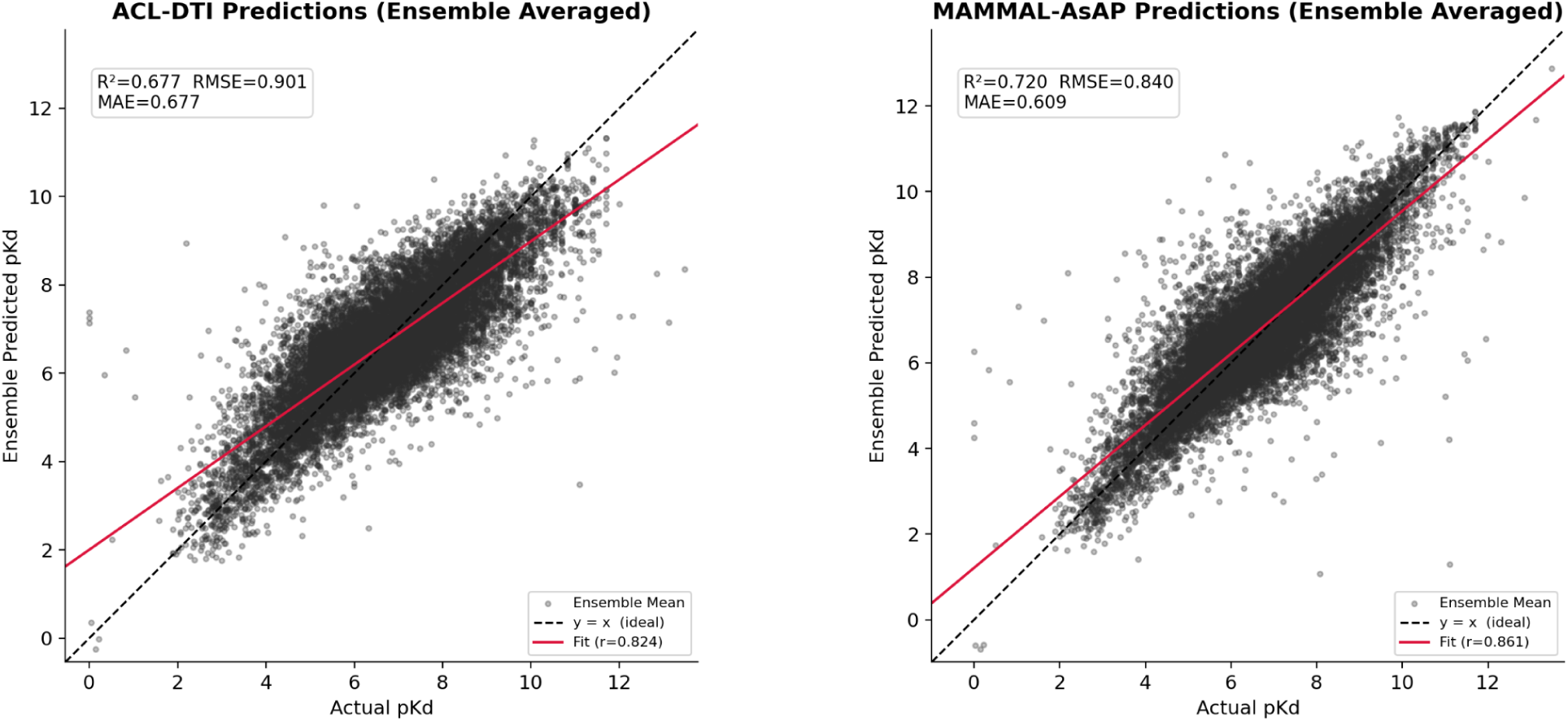
The ensemble-averaged predictions from ACL and MAMMAL-AsAP show similar correspondence with the measured (pK_d_) values, with (R^2^) values of (0.677) and (0.720), respectively. The predicted values span the observed affinity range for both models and show a consistent positive relationship with the measurements. The fitted correlations are also similar (r=0.824 for ACL and r=0.86 for MAMMAL-AsAP), indicating comparable agreement between predicted and measured affinities. Together, these results suggest that both approaches capture the major variation in binding affinity without a substantial difference in overall predictive behavior.

## Supplementary Tables

### Benchmark Performance of Assay-Aware and Assay-Agnostic Models Across Random Seeds

**Table 2:**
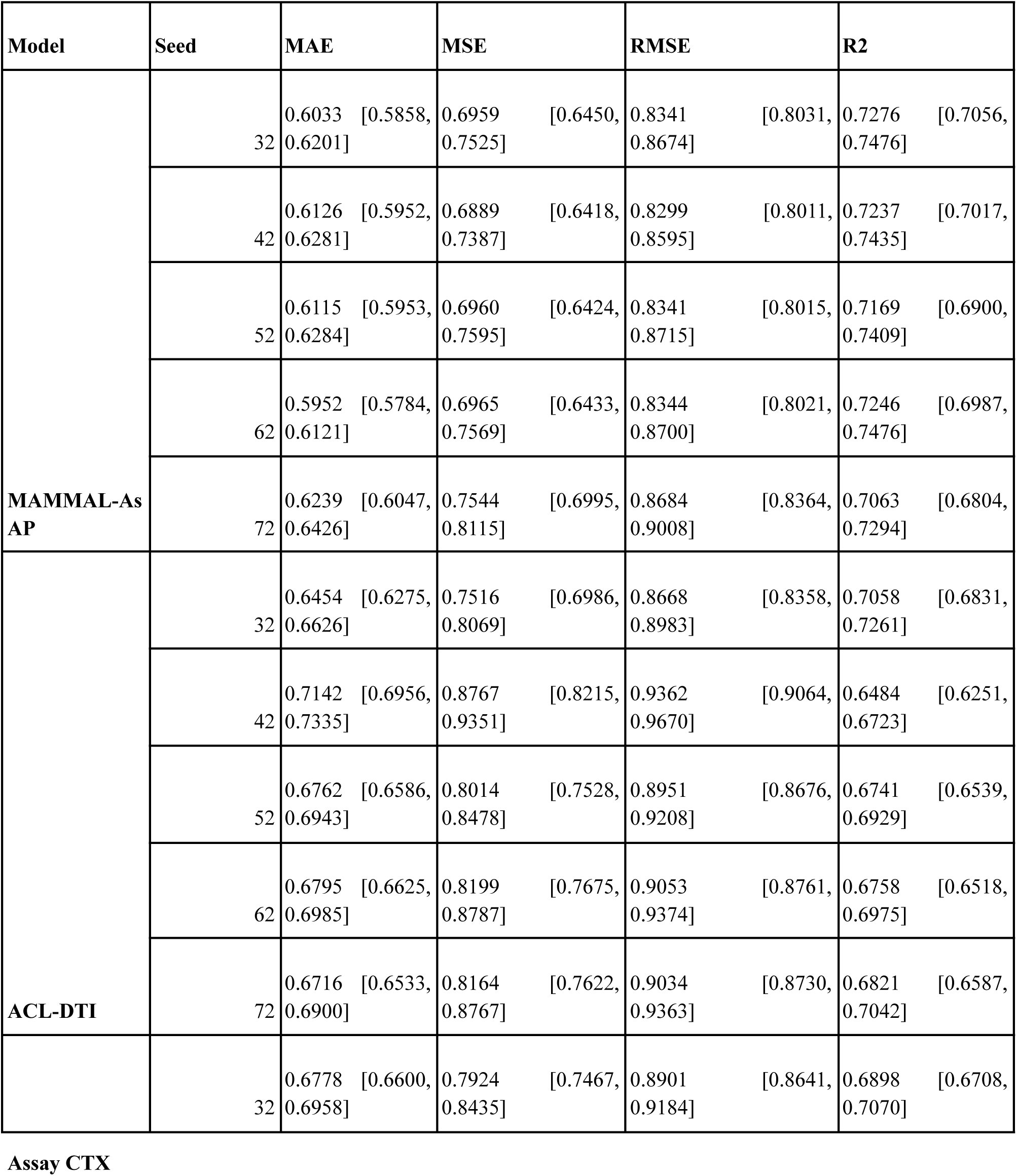
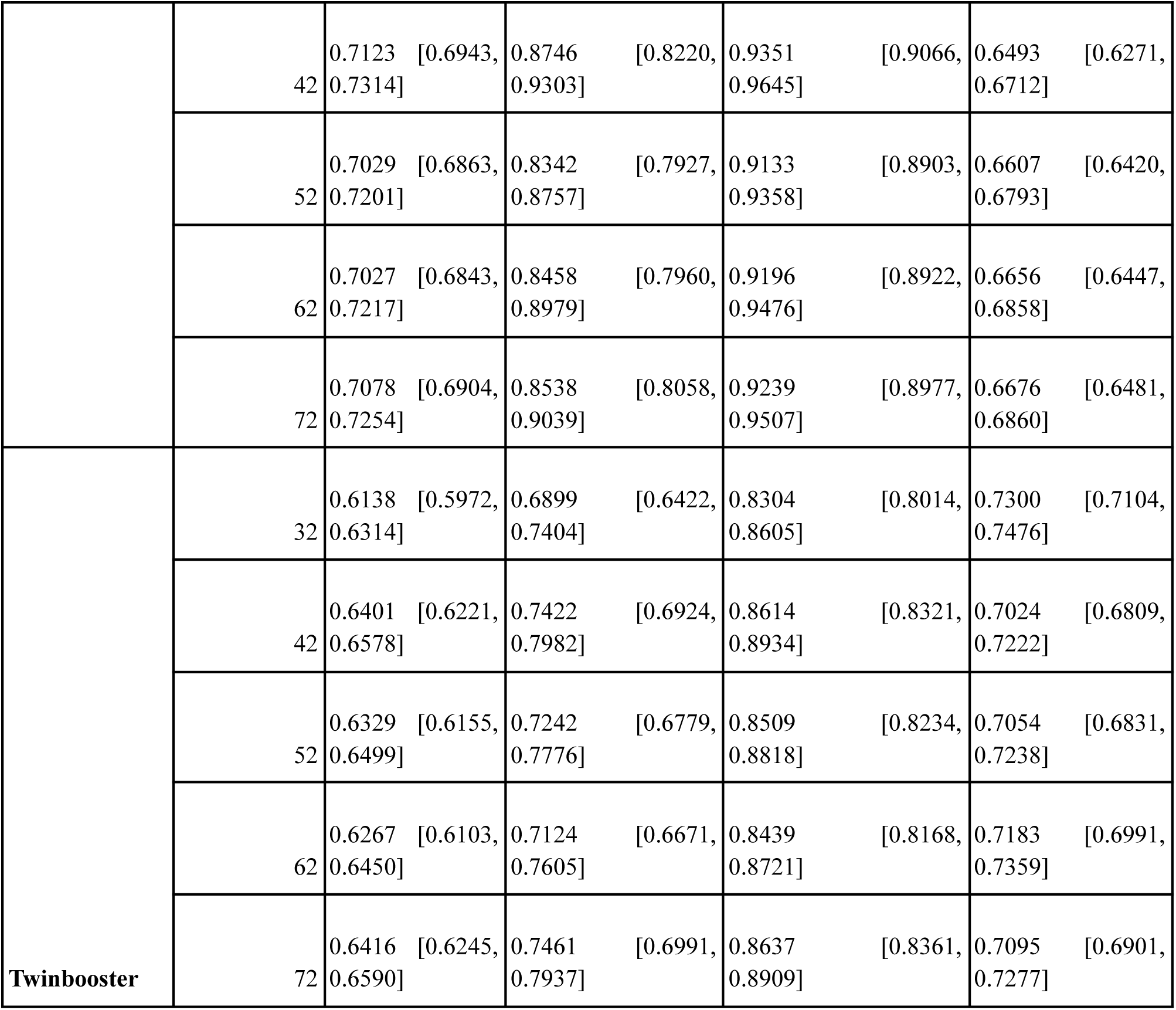
Performance of assay-aware models across five random seeds.

| Model | Seed | MAE | MSE | RMSE | R2 |
| --- | --- | --- | --- | --- | --- |
| MAMMAL-As AP | 32 | 0.6033 [0.5858, 0.6201] | 0.6959 [0.6450, 0.7525] | 0.8341 [0.8031, 0.8674] | 0.7276 [0.7056, 0.7476] |
|  | 42 | 0.6126 [0.5952, 0.6281] | 0.6889 [0.6418, 0.7387] | 0.8299 [0.8011, 0.8595] | 0.7237 [0.7017, 0.7435] |
|  | 52 | 0.6115 [0.5953, 0.6284] | 0.6960 [0.6424, 0.7595] | 0.8341 [0.8015, 0.8715] | 0.7169 [0.6900, 0.7409] |
|  | 62 | 0.5952 [0.5784, 0.6121] | 0.6965 [0.6433, 0.7569] | 0.8344 [0.8021, 0.8700] | 0.7246 [0.6987, 0.7476] |
|  | 72 | 0.6239 [0.6047, 0.6426] | 0.7544 [0.6995, 0.8115] | 0.8684 [0.8364, 0.9008] | 0.7063 [0.6804, 0.7294] |
| ACL-DTI | 32 | 0.6454 [0.6275, 0.6626] | 0.7516 [0.6986, 0.8069] | 0.8668 [0.8358, 0.8983] | 0.7058 [0.6831, 0.7261] |
|  | 42 | 0.7142 [0.6956, 0.7335] | 0.8767 [0.8215, 0.9351] | 0.9362 [0.9064, 0.9670] | 0.6484 [0.6251, 0.6723] |
|  | 52 | 0.6762 [0.6586, 0.6943] | 0.8014 [0.7528, 0.8478] | 0.8951 [0.8676, 0.9208] | 0.6741 [0.6539, 0.6929] |
|  | 62 | 0.6795 [0.6625, 0.6985] | 0.8199 [0.7675, 0.8787] | 0.9053 [0.8761, 0.9374] | 0.6758 [0.6518, 0.6975] |
|  | 72 | 0.6716 [0.6533, 0.6900] | 0.8164 [0.7622, 0.8767] | 0.9034 [0.8730, 0.9363] | 0.6821 [0.6587, 0.7042] |
|  | 32 | 0.6778 [0.6600, 0.6958] | 0.7924 [0.7467, 0.8435] | 0.8901 [0.8641, 0.9184] | 0.6898 [0.6708, 0.7070] |
|  | 42 | 0.7123 [0.6943, 0.7314] | 0.8746 [0.8220, 0.9303] | 0.9351 [0.9066, 0.9645] | 0.6493 [0.6271, 0.6712] |
|  | 52 | 0.7029 [0.6863, 0.7201] | 0.8342 [0.7927, 0.8757] | 0.9133 [0.8903, 0.9358] | 0.6607 [0.6420, 0.6793] |
|  | 62 | 0.7027 [0.6843, 0.7217] | 0.8458 [0.7960, 0.8979] | 0.9196 [0.8922, 0.9476] | 0.6656 [0.6447, 0.6858] |
|  | 72 | 0.7078 [0.6904, 0.7254] | 0.8538 [0.8058, 0.9039] | 0.9239 [0.8977, 0.9507] | 0.6676 [0.6481, 0.6860] |
| **Twinbooster** | 32 | 0.6138 [0.5972, 0.6314] | 0.6899 [0.6422, 0.7404] | 0.8304 [0.8014, 0.8605] | 0.7300 [0.7104, 0.7476] |
|  | 42 | 0.6401 [0.6221, 0.6578] | 0.7422 [0.6924, 0.7982] | 0.8614 [0.8321, 0.8934] | 0.7024 [0.6809, 0.7222] |
|  | 52 | 0.6329 [0.6155, 0.6499] | 0.7242 [0.6779, 0.7776] | 0.8509 [0.8234, 0.8818] | 0.7054 [0.6831, 0.7238] |
|  | 62 | 0.6267 [0.6103, 0.6450] | 0.7124 [0.6671, 0.7605] | 0.8439 [0.8168, 0.8721] | 0.7183 [0.6991, 0.7359] |
|  | 72 | 0.6416 [0.6245, 0.6590] | 0.7461 [0.6991, 0.7937] | 0.8637 [0.8361, 0.8909] | 0.7095 [0.6901, 0.7277] |

**Table 3:**
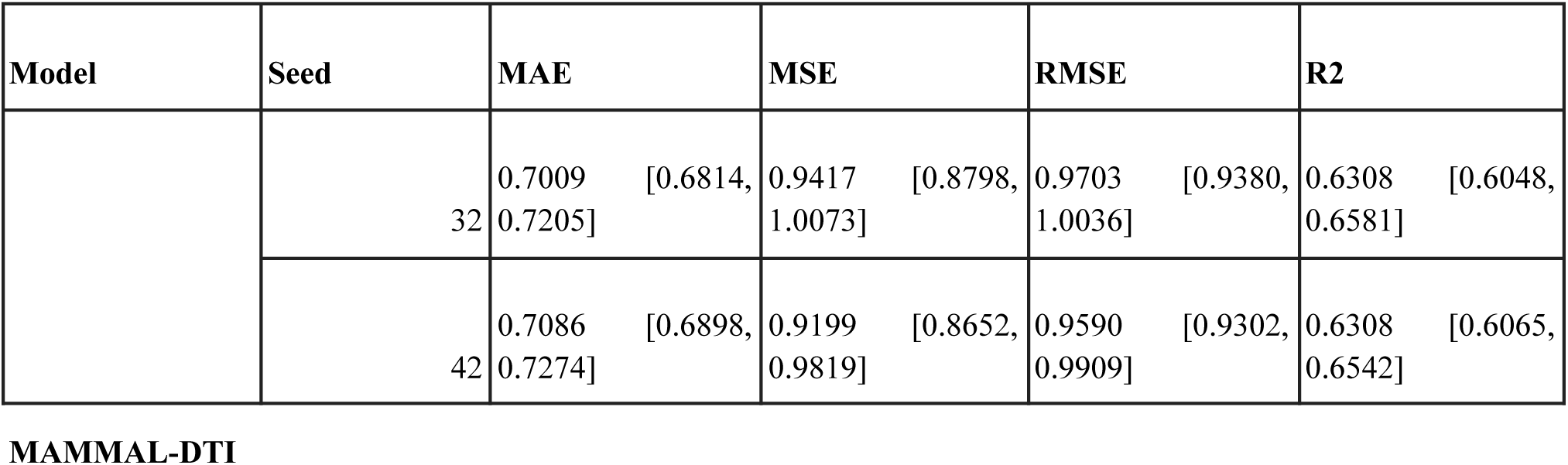
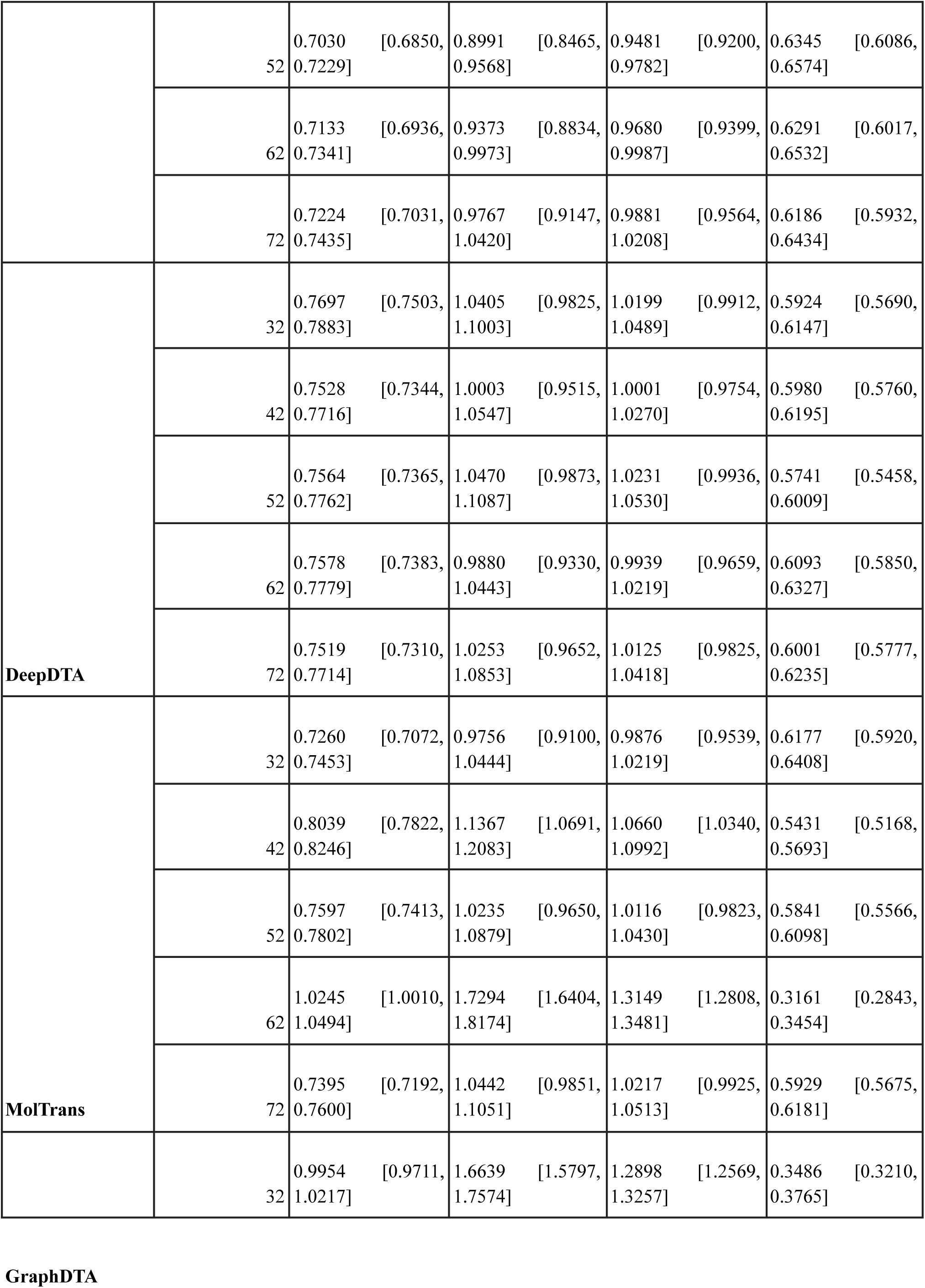
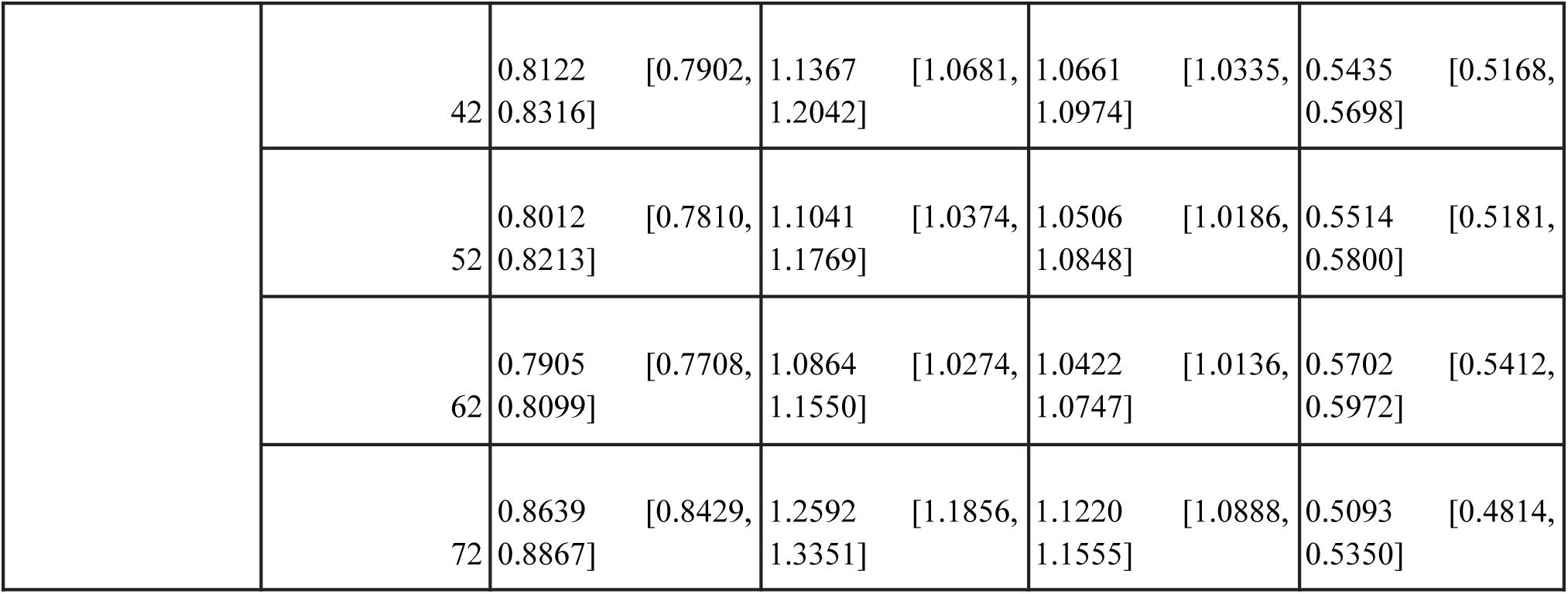
Performance of assay-agnostic models across five random seeds.

| Model | Seed | MAE | MSE | RMSE | R2 |
| --- | --- | --- | --- | --- | --- |
|  | 32 | 0.7009 [0.6814, 0.7205] | 0.9417 [0.8798, 1.0073] | 0.9703 [0.9380, 1.0036] | 0.6308 [0.6048, 0.6581] |
|  | 42 | 0.7086 [0.6898, 0.7274] | 0.9199 [0.8652, 0.9819] | 0.9590 [0.9302, 0.9909] | 0.6308 [0.6065, 0.6542] |
| **MAMMAL-DTI** |  |  |  |  |  |
|  | 52 | 0.7030 [0.6850, 0.7229] | 0.8991 [0.8465, 0.9568] | 0.9481 [0.9200, 0.9782] | 0.6345 [0.6086, 0.6574] |
|  | 62 | 0.7133 [0.6936, 0.7341] | 0.9373 [0.8834, 0.9973] | 0.9680 [0.9399, 0.9987] | 0.6291 [0.6017, 0.6532] |
|  | 72 | 0.7224 [0.7031, 0.7435] | 0.9767 [0.9147, 1.0420] | 0.9881 [0.9564, 1.0208] | 0.6186 [0.5932, 0.6434] |
| **DeepDTA** | 32 | 0.7697 [0.7503, 0.7883] | 1.0405 [0.9825, 1.1003] | 1.0199 [0.9912, 1.0489] | 0.5924 [0.5690, 0.6147] |
|  | 42 | 0.7528 [0.7344, 0.7716] | 1.0003 [0.9515, 1.0547] | 1.0001 [0.9754, 1.0270] | 0.5980 [0.5760, 0.6195] |
|  | 52 | 0.7564 [0.7365, 0.7762] | 1.0470 [0.9873, 1.1087] | 1.0231 [0.9936, 1.0530] | 0.5741 [0.5458, 0.6009] |
|  | 62 | 0.7578 [0.7383, 0.7779] | 0.9880 [0.9330, 1.0443] | 0.9939 [0.9659, 1.0219] | 0.6093 [0.5850, 0.6327] |
|  | 72 | 0.7519 [0.7310, 0.7714] | 1.0253 [0.9652, 1.0853] | 1.0125 [0.9825, 1.0418] | 0.6001 [0.5777, 0.6235] |
| **MolTrans** | 32 | 0.7260 [0.7072, 0.7453] | 0.9756 [0.9100, 1.0444] | 0.9876 [0.9539, 1.0219] | 0.6177 [0.5920, 0.6408] |
|  | 42 | 0.8039 [0.7822, 0.8246] | 1.1367 [1.0691, 1.2083] | 1.0660 [1.0340, 1.0992] | 0.5431 [0.5168, 0.5693] |
|  | 52 | 0.7597 [0.7413, 0.7802] | 1.0235 [0.9650, 1.0879] | 1.0116 [0.9823, 1.0430] | 0.5841 [0.5566, 0.6098] |
|  | 62 | 1.0245 [1.0010, 1.0494] | 1.7294 [1.6404, 1.8174] | 1.3149 [1.2808, 1.3481] | 0.3161 [0.2843, 0.3454] |
|  | 72 | 0.7395 [0.7192, 0.7600] | 1.0442 [0.9851, 1.1051] | 1.0217 [0.9925, 1.0513] | 0.5929 [0.5675, 0.6181] |
|  | 32 | 0.9954 [0.9711, 1.0217] | 1.6639 [1.5797, 1.7574] | 1.2898 [1.2569, 1.3257] | 0.3486 [0.3210, 0.3765] |
|  | 42 | 0.8122 [0.7902, 0.8316] | 1.1367 [1.0681, 1.2042] | 1.0661 [1.0335, 1.0974] | 0.5435 [0.5168, 0.5698] |
|  | 52 | 0.8012 [0.7810, 0.8213] | 1.1041 [1.0374, 1.1769] | 1.0506 [1.0186, 1.0848] | 0.5514 [0.5181, 0.5800] |
|  | 62 | 0.7905 [0.7708, 0.8099] | 1.0864 [1.0274, 1.1550] | 1.0422 [1.0136, 1.0747] | 0.5702 [0.5412, 0.5972] |
|  | 72 | 0.8639 [0.8429, 0.8867] | 1.2592 [1.1856, 1.3351] | 1.1220 [1.0888, 1.1555] | 0.5093 [0.4814, 0.5350] |

**Table 4:** Mean performance of assay-agnostic models across five random seeds. Values are reported as mean ± standard deviation

| Model | MAE | MSE | RMSE | R2 |
| --- | --- | --- | --- | --- |
| DeepDTA | 0.7577 ± 0.0071 | 1.0202 ± 0.0254 | 1.0099 ± 0.0126 | 0.5948 ± 0.0131 |
| GraphDTA | 0.8526 ± 0.0846 | 1.2501 ± 0.2410 | 1.1141 ± 0.1030 | 0.5046 ± 0.0900 |
| MAMMAL-AsAP | 0.7096 ± 0.0086 | 0.9349 ± 0.0287 | 0.9667 ± 0.0148 | 0.6288 ± 0.0060 |
| MolTrans | 0.8107 ± 0.1231 | 1.1819 ± 0.3116 | 1.0804 ± 0.1342 | 0.5308 ± 0.1230 |

### Benchmark Results Under Assay-Description Masking

**Table 5:**
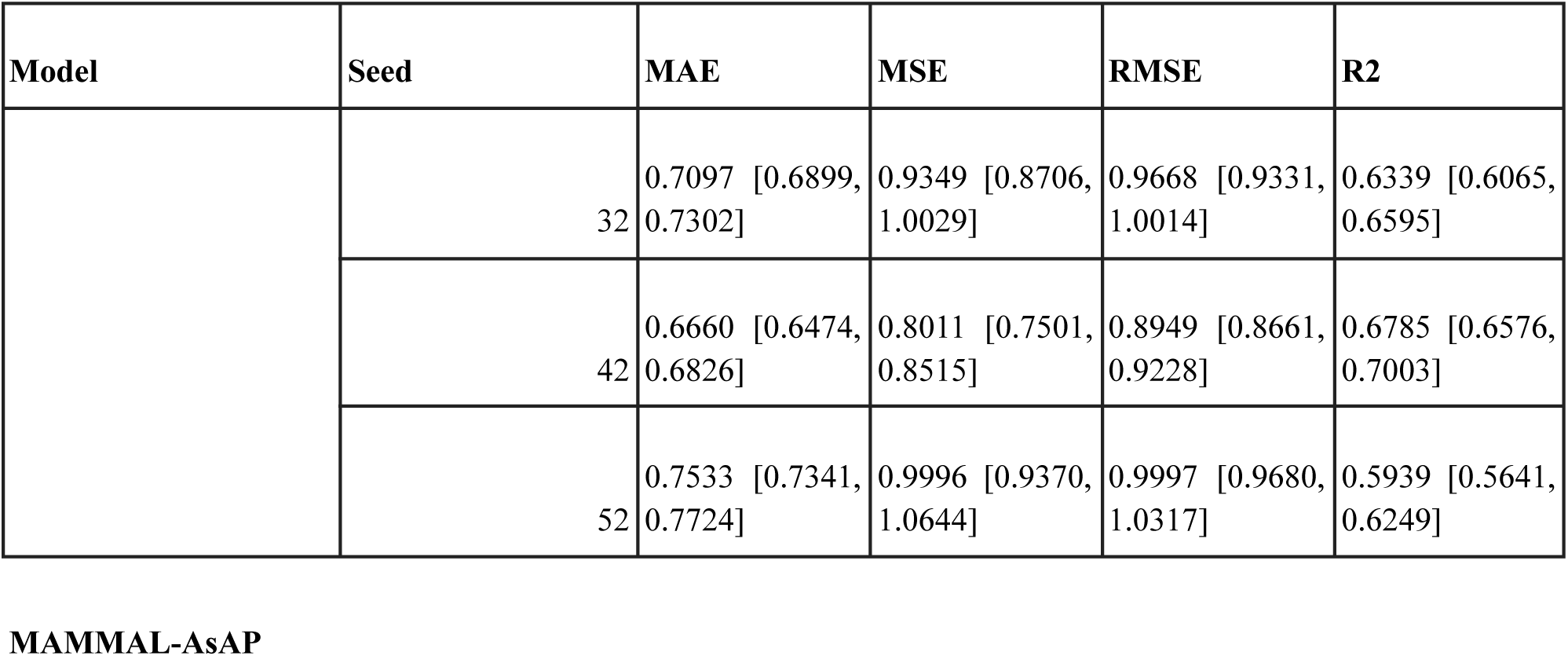
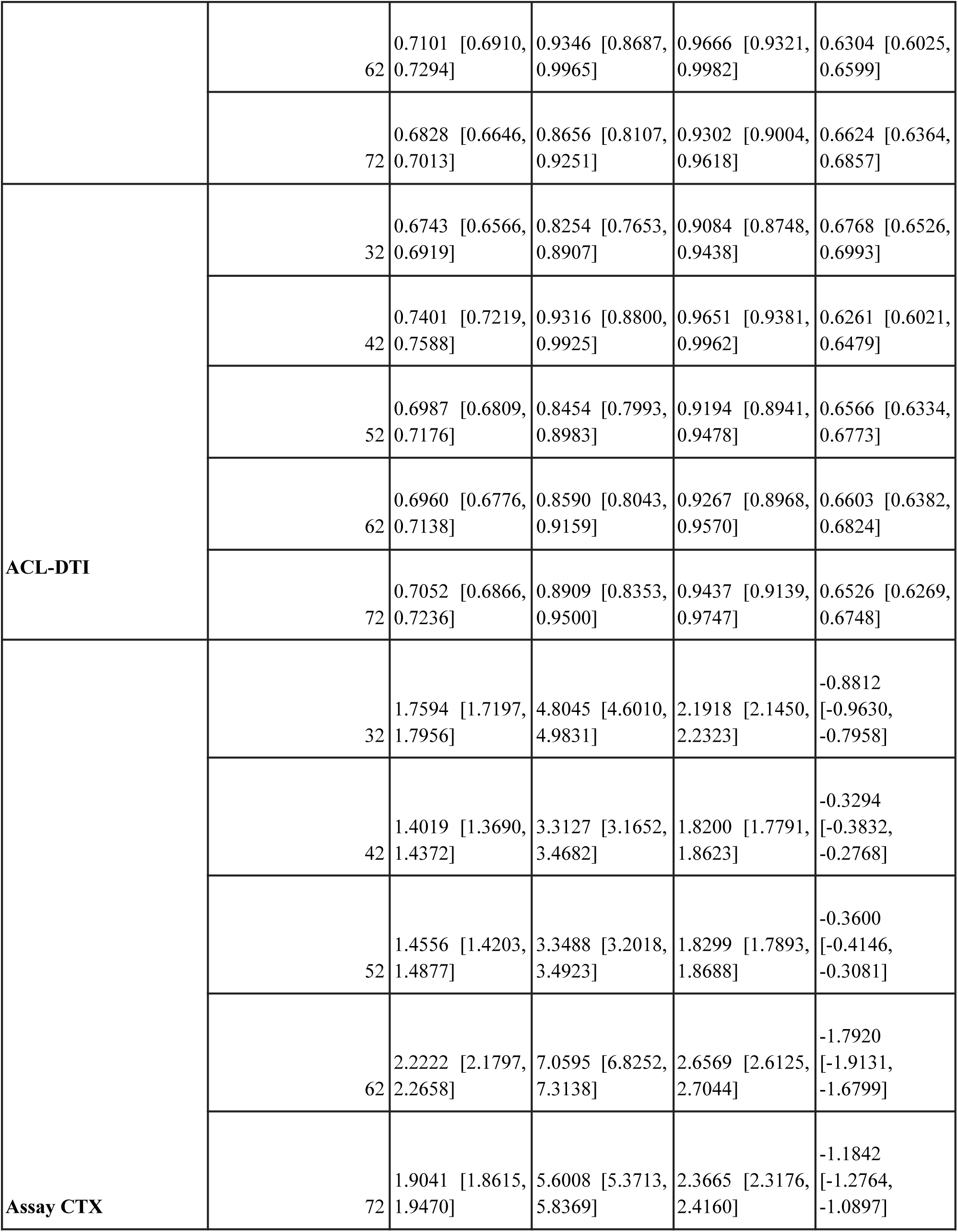
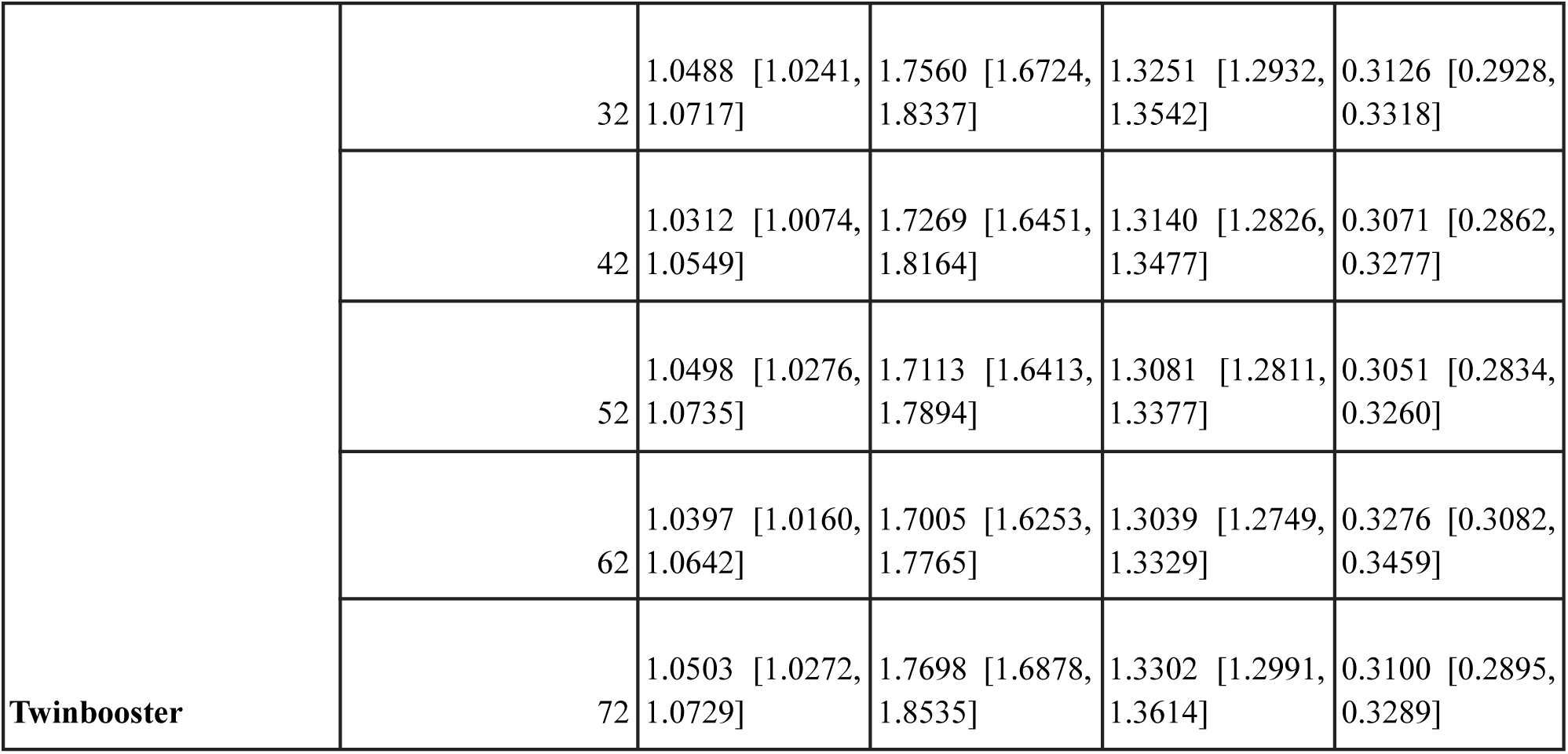
Performance evaluation on test sets with target names masked from assay descriptions across five random seeds.

| Model | Seed | MAE | MSE | RMSE | R2 |
| --- | --- | --- | --- | --- | --- |
|  | 32 | 0.7097 [0.6899, 0.7302] | 0.9349 [0.8706, 1.0029] | 0.9668 [0.9331, 1.0014] | 0.6339 [0.6065, 0.6595] |
|  | 42 | 0.6660 [0.6474, 0.6826] | 0.8011 [0.7501, 0.8515] | 0.8949 [0.8661, 0.9228] | 0.6785 [0.6576, 0.7003] |
|  | 52 | 0.7533 [0.7341, 0.7724] | 0.9996 [0.9370, 1.0644] | 0.9997 [0.9680, 1.0317] | 0.5939 [0.5641, 0.6249] |
|  | 62 | 0.7101 [0.6910, 0.7294] | 0.9346 [0.8687, 0.9965] | 0.9666 [0.9321, 0.9982] | 0.6304 [0.6025, 0.6599] |
|  | 72 | 0.6828 [0.6646, 0.7013] | 0.8656 [0.8107, 0.9251] | 0.9302 [0.9004, 0.9618] | 0.6624 [0.6364, 0.6857] |
| ACL-DTI | 32 | 0.6743 [0.6566, 0.6919] | 0.8254 [0.7653, 0.8907] | 0.9084 [0.8748, 0.9438] | 0.6768 [0.6526, 0.6993] |
|  | 42 | 0.7401 [0.7219, 0.7588] | 0.9316 [0.8800, 0.9925] | 0.9651 [0.9381, 0.9962] | 0.6261 [0.6021, 0.6479] |
|  | 52 | 0.6987 [0.6809, 0.7176] | 0.8454 [0.7993, 0.8983] | 0.9194 [0.8941, 0.9478] | 0.6566 [0.6334, 0.6773] |
|  | 62 | 0.6960 [0.6776, 0.7138] | 0.8590 [0.8043, 0.9159] | 0.9267 [0.8968, 0.9570] | 0.6603 [0.6382, 0.6824] |
|  | 72 | 0.7052 [0.6866, 0.7236] | 0.8909 [0.8353, 0.9500] | 0.9437 [0.9139, 0.9747] | 0.6526 [0.6269, 0.6748] |
| Assay CTX | 32 | 1.7594 [1.7197, 1.7956] | 4.8045 [4.6010, 4.9831] | 2.1918 [2.1450, 2.2323] | -0.8812 [-0.9630, -0.7958] |
|  | 42 | 1.4019 [1.3690, 1.4372] | 3.3127 [3.1652, 3.4682] | 1.8200 [1.7791, 1.8623] | -0.3294 [-0.3832, -0.2768] |
|  | 52 | 1.4556 [1.4203, 1.4877] | 3.3488 [3.2018, 3.4923] | 1.8299 [1.7893, 1.8688] | -0.3600 [-0.4146, -0.3081] |
|  | 62 | 2.2222 [2.1797, 2.2658] | 7.0595 [6.8252, 7.3138] | 2.6569 [2.6125, 2.7044] | -1.7920 [-1.9131, -1.6799] |
|  | 72 | 1.9041 [1.8615, 1.9470] | 5.6008 [5.3713, 5.8369] | 2.3665 [2.3176, 2.4160] | -1.1842 [-1.2764, -1.0897] |
| **Twinbooster** | 32 | 1.0488 [1.0241, 1.0717] | 1.7560 [1.6724, 1.8337] | 1.3251 [1.2932, 1.3542] | 0.3126 [0.2928, 0.3318] |
|  | 42 | 1.0312 [1.0074, 1.0549] | 1.7269 [1.6451, 1.8164] | 1.3140 [1.2826, 1.3477] | 0.3071 [0.2862, 0.3277] |
|  | 52 | 1.0498 [1.0276, 1.0735] | 1.7113 [1.6413, 1.7894] | 1.3081 [1.2811, 1.3377] | 0.3051 [0.2834, 0.3260] |
|  | 62 | 1.0397 [1.0160, 1.0642] | 1.7005 [1.6253, 1.7765] | 1.3039 [1.2749, 1.3329] | 0.3276 [0.3082, 0.3459] |
|  | 72 | 1.0503 [1.0272, 1.0729] | 1.7698 [1.6878, 1.8535] | 1.3302 [1.2991, 1.3614] | 0.3100 [0.2895, 0.3289] |

**Table 6:** Average performance of all models on test sets with target names masked. Values are reported as mean ± standard deviation across five random seeds.

| **Model** | **MAE** | **MSE** | **RMSE** | **R2** |
| --- | --- | --- | --- | --- |
| **Assay CTX** | 1.749 ± 0.337 | 4.825 ± 1.586 | 2.173 ± 0.359 | -0.909 ± 0.611 |
| **ACL-DTI** | 0.703 ± 0.024 | 0.870 ± 0.042 | 0.933 ± 0.022 | 0.654 ± 0.018 |
| **MAMMAL-AsAP** | 0.704 ± 0.033 | 0.907 ± 0.076 | 0.952 ± 0.040 | 0.640 ± 0.033 |
| **Twinbooster** | 1.044 ± 0.008 | 1.733 ± 0.029 | 1.316 ± 0.011 | 0.312 ± 0.009 |

**Table 7:**
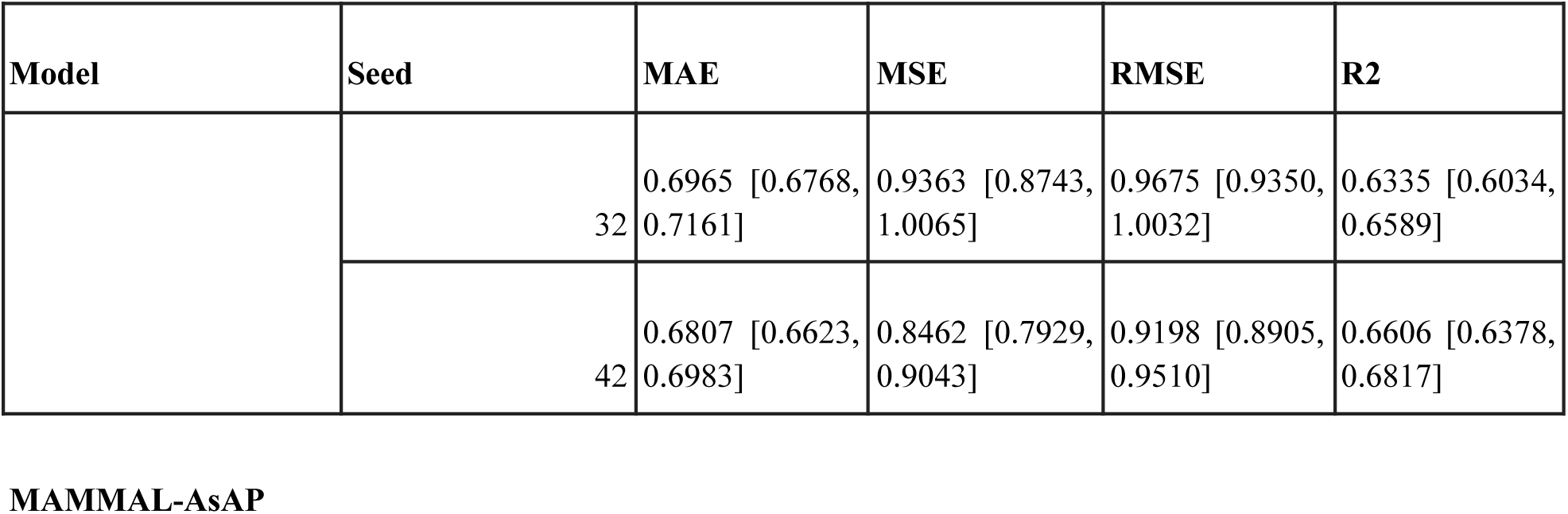
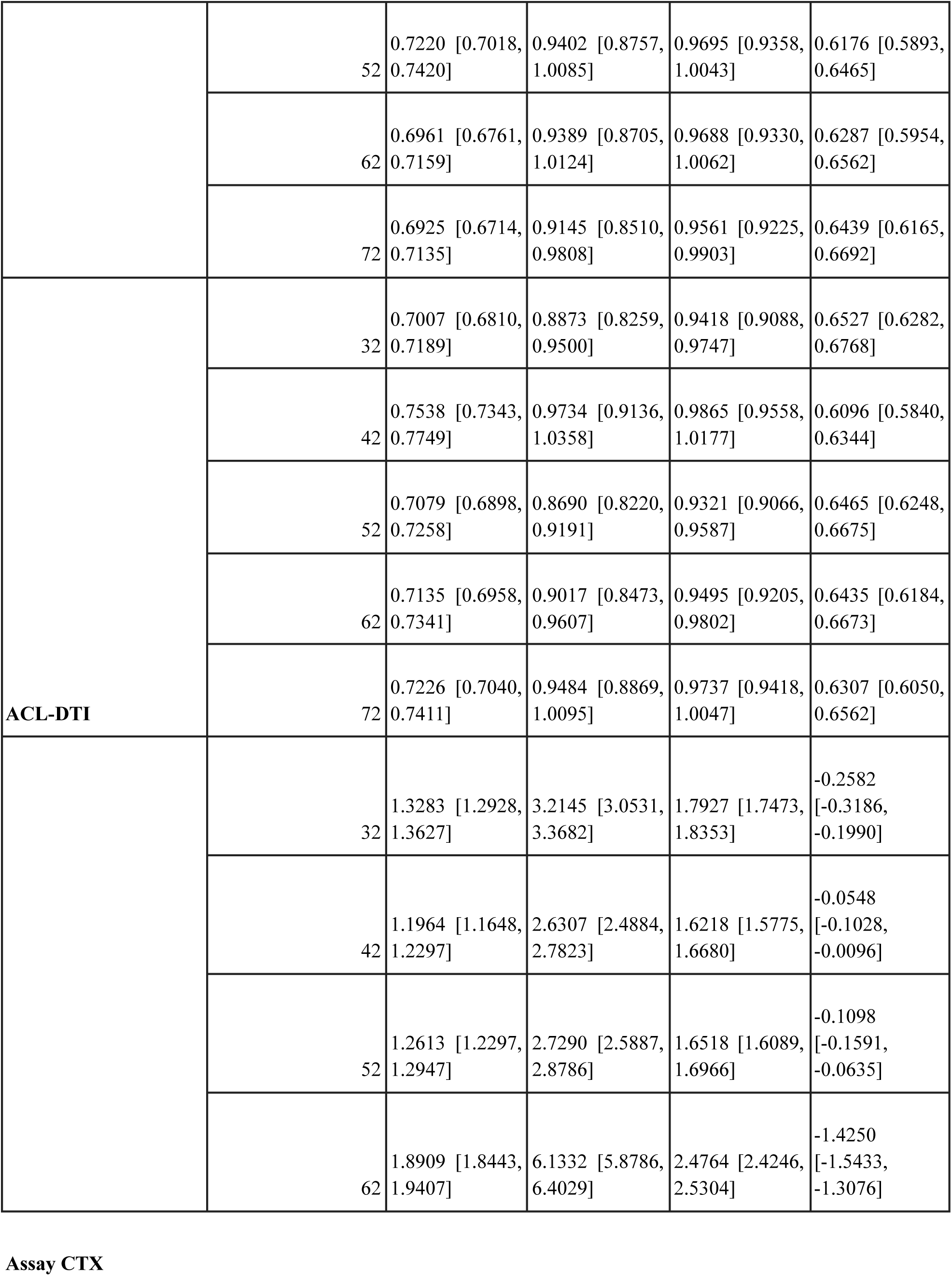
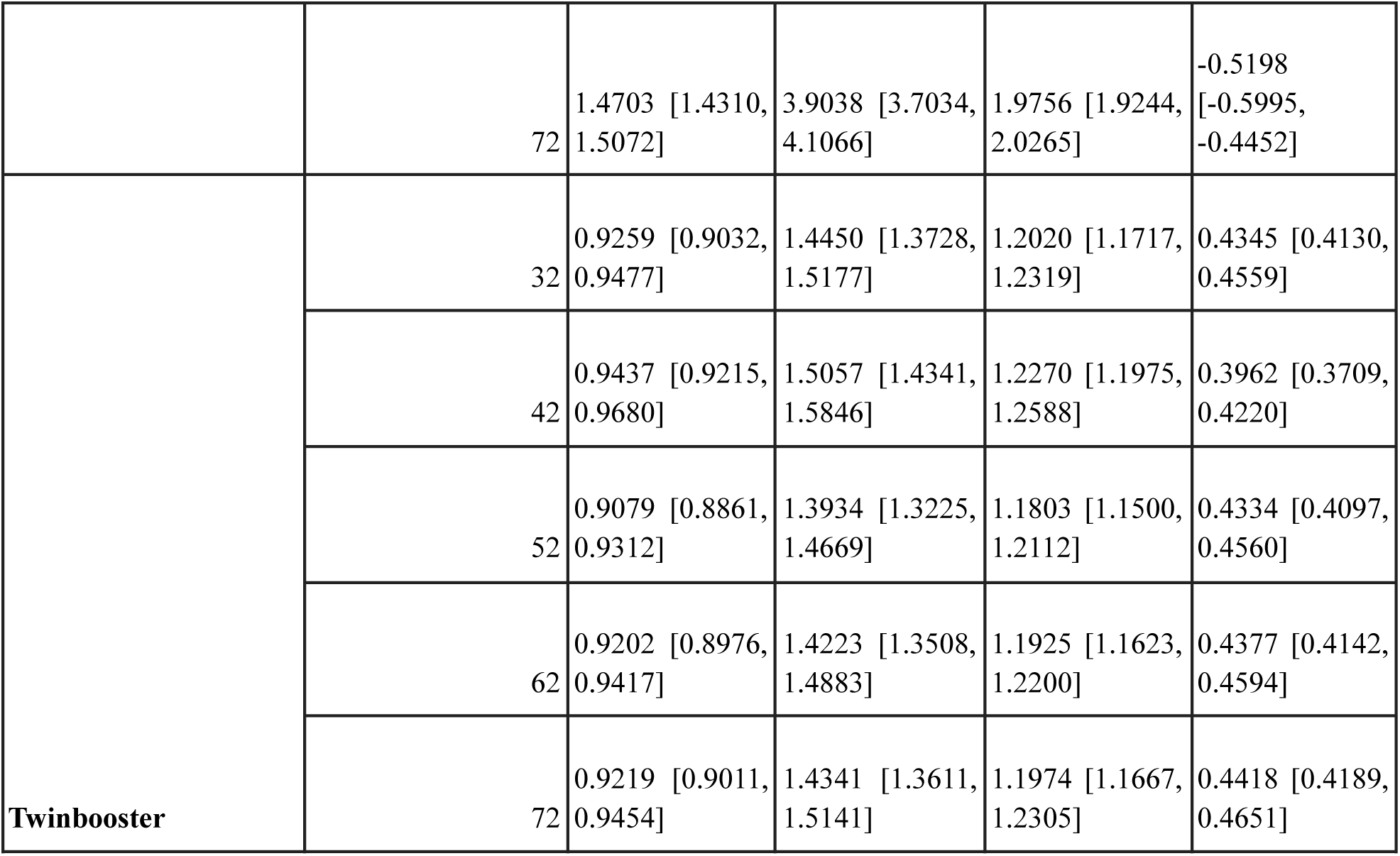
Performance evaluation on test sets with assay techniques masked from assay descriptions across five random seeds.

**Table 8:** Average performance of all models on test sets with assay techniques masked. Values are reported as mean ± standard deviation across five random seeds.

| **Model** | **MAE** | **MSE** | **RMSE** | **R2** |
| --- | --- | --- | --- | --- |
| **Assay CTX** | 1.429 ± 0.277 | 3.722 ± 1.439 | 1.904 ± 0.349 | -0.474 ± 0.562 |
| **ACL-DTI** | 0.720 ± 0.021 | 0.916 ± 0.044 | 0.957 ± 0.023 | 0.637 ± 0.017 |
| **MAMMAL-AsAP** | 0.698 ± 0.015 | 0.915 ± 0.040 | 0.956 ± 0.021 | 0.637 ± 0.016 |
| **Twinbooster** | 0.924 ± 0.013 | 1.440 ± 0.041 | 1.200 ± 0.017 | 0.429 ± 0.018 |

**Table 9:**
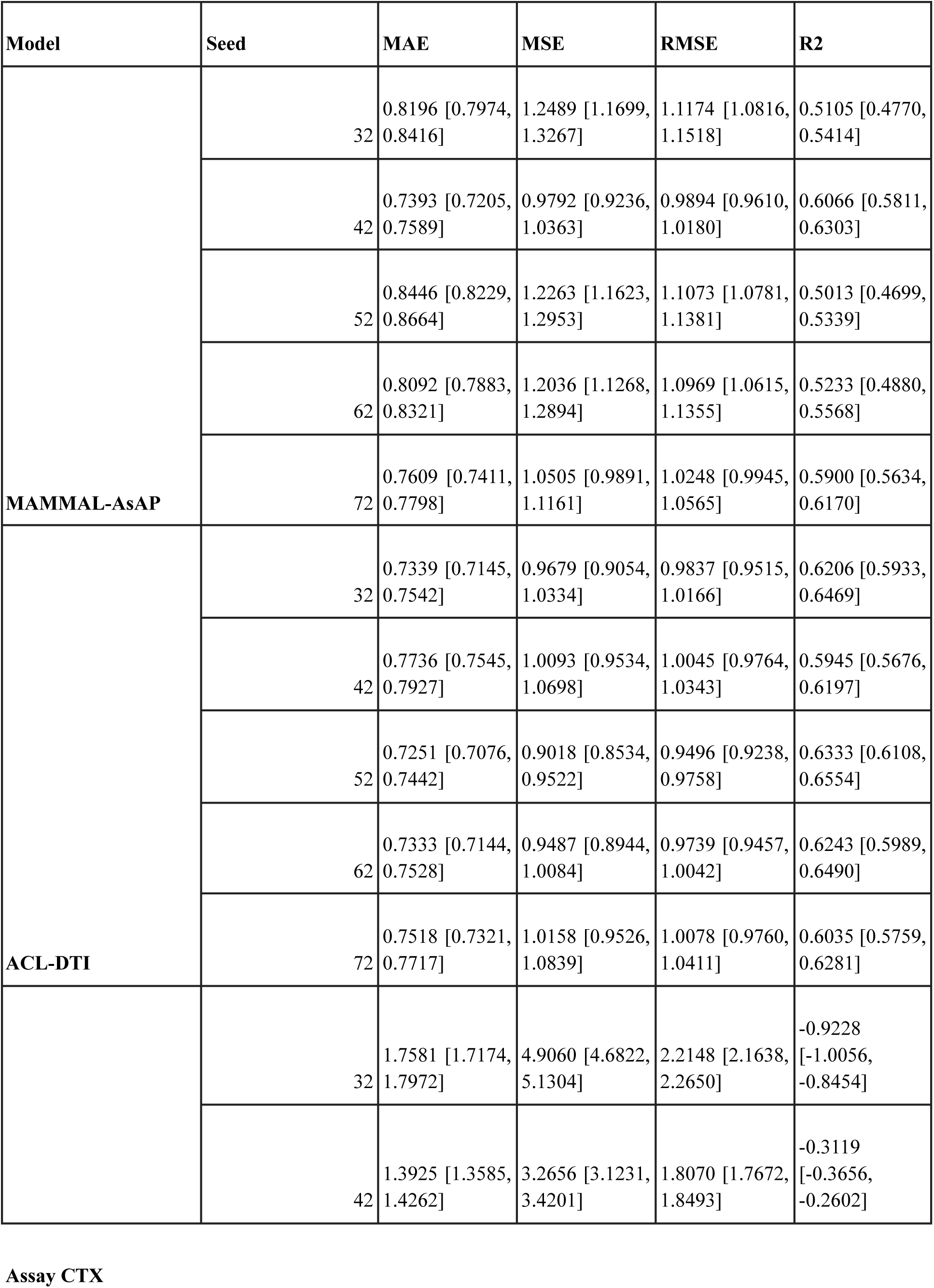
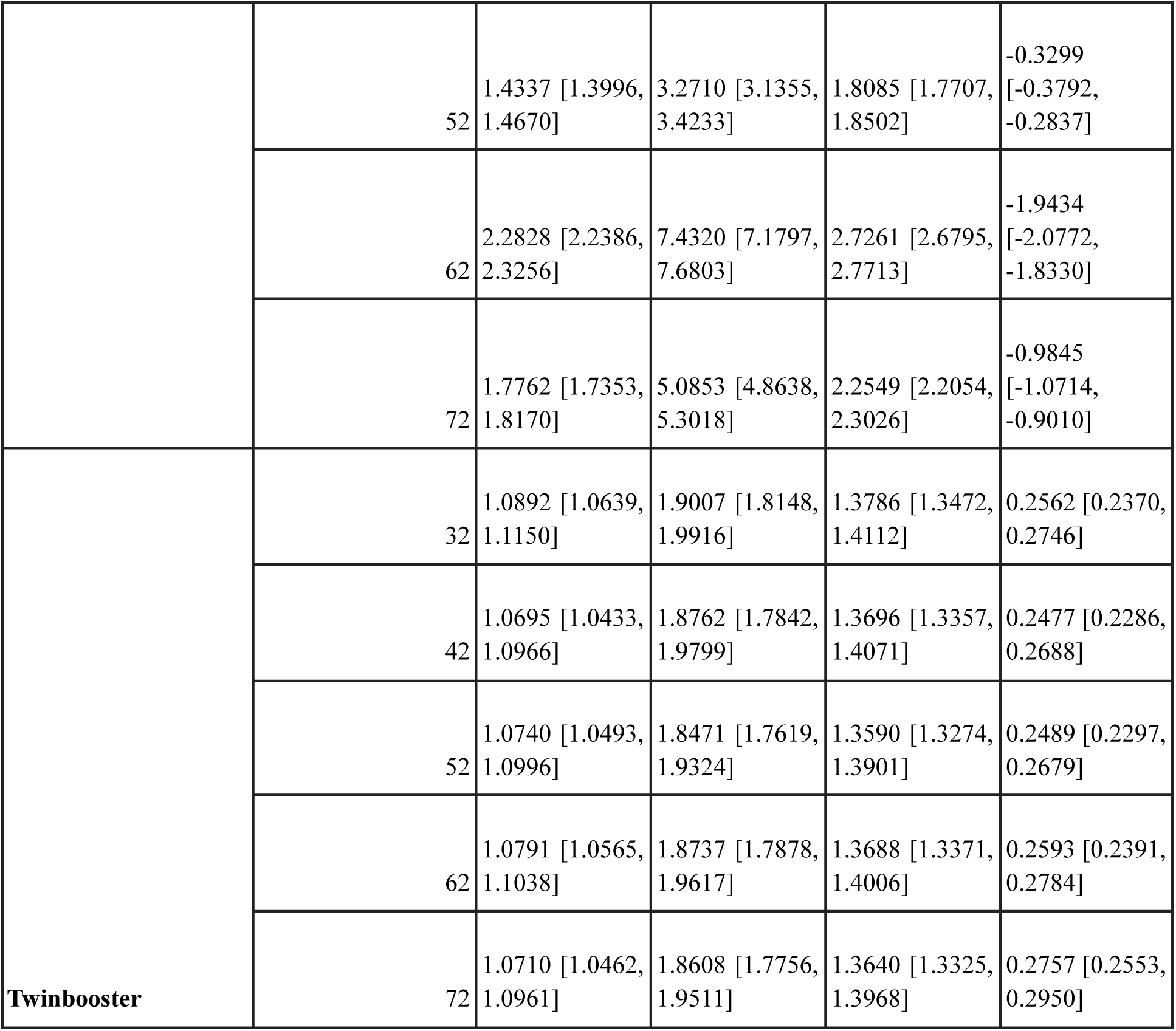
Performance evaluation on test sets with assay techniques and target names both masked from assay descriptions across five random seeds.

| Model | Seed | MAE | MSE | RMSE | R2 |
| --- | --- | --- | --- | --- | --- |
| MAMMAL-AsAP | 32 | 0.8196 [0.7974, 0.8416] | 1.2489 [1.1699, 1.3267] | 1.1174 [1.0816, 1.1518] | 0.5105 [0.4770, 0.5414] |
|  | 42 | 0.7393 [0.7205, 0.7589] | 0.9792 [0.9236, 1.0363] | 0.9894 [0.9610, 1.0180] | 0.6066 [0.5811, 0.6303] |
|  | 52 | 0.8446 [0.8229, 0.8664] | 1.2263 [1.1623, 1.2953] | 1.1073 [1.0781, 1.1381] | 0.5013 [0.4699, 0.5339] |
|  | 62 | 0.8092 [0.7883, 0.8321] | 1.2036 [1.1268, 1.2894] | 1.0969 [1.0615, 1.1355] | 0.5233 [0.4880, 0.5568] |
|  | 72 | 0.7609 [0.7411, 0.7798] | 1.0505 [0.9891, 1.1161] | 1.0248 [0.9945, 1.0565] | 0.5900 [0.5634, 0.6170] |
| ACL-DTI | 32 | 0.7339 [0.7145, 0.7542] | 0.9679 [0.9054, 1.0334] | 0.9837 [0.9515, 1.0166] | 0.6206 [0.5933, 0.6469] |
|  | 42 | 0.7736 [0.7545, 0.7927] | 1.0093 [0.9534, 1.0698] | 1.0045 [0.9764, 1.0343] | 0.5945 [0.5676, 0.6197] |
|  | 52 | 0.7251 [0.7076, 0.7442] | 0.9018 [0.8534, 0.9522] | 0.9496 [0.9238, 0.9758] | 0.6333 [0.6108, 0.6554] |
|  | 62 | 0.7333 [0.7144, 0.7528] | 0.9487 [0.8944, 1.0084] | 0.9739 [0.9457, 1.0042] | 0.6243 [0.5989, 0.6490] |
|  | 72 | 0.7518 [0.7321, 0.7717] | 1.0158 [0.9526, 1.0839] | 1.0078 [0.9760, 1.0411] | 0.6035 [0.5759, 0.6281] |
|  | 32 | 1.7581 [1.7174, 1.7972] | 4.9060 [4.6822, 5.1304] | 2.2148 [2.1638, 2.2650] | -0.9228 [-1.0056, -0.8454] |
|  | 42 | 1.3925 [1.3585, 1.4262] | 3.2656 [3.1231, 3.4201] | 1.8070 [1.7672, 1.8493] | -0.3119 [-0.3656, -0.2602] |
|  | 52 | 1.4337 [1.3996, 1.4670] | 3.2710 [3.1355, 3.4233] | 1.8085 [1.7707, 1.8502] | -0.3299 [-0.3792, -0.2837] |
|  | 62 | 2.2828 [2.2386, 2.3256] | 7.4320 [7.1797, 7.6803] | 2.7261 [2.6795, 2.7713] | -1.9434 [-2.0772, -1.8330] |
|  | 72 | 1.7762 [1.7353, 1.8170] | 5.0853 [4.8638, 5.3018] | 2.2549 [2.2054, 2.3026] | -0.9845 [-1.0714, -0.9010] |
| **Twinbooster** | 32 | 1.0892 [1.0639, 1.1150] | 1.9007 [1.8148, 1.9916] | 1.3786 [1.3472, 1.4112] | 0.2562 [0.2370, 0.2746] |
|  | 42 | 1.0695 [1.0433, 1.0966] | 1.8762 [1.7842, 1.9799] | 1.3696 [1.3357, 1.4071] | 0.2477 [0.2286, 0.2688] |
|  | 52 | 1.0740 [1.0493, 1.0996] | 1.8471 [1.7619, 1.9324] | 1.3590 [1.3274, 1.3901] | 0.2489 [0.2297, 0.2679] |
|  | 62 | 1.0791 [1.0565, 1.1038] | 1.8737 [1.7878, 1.9617] | 1.3688 [1.3371, 1.4006] | 0.2593 [0.2391, 0.2784] |
|  | 72 | 1.0710 [1.0462, 1.0961] | 1.8608 [1.7756, 1.9511] | 1.3640 [1.3325, 1.3968] | 0.2757 [0.2553, 0.2950] |

**Table 10:** Average performance of all models on test sets with assay techniques and target names both masked. Values are reported as mean ± standard deviation across five random seeds.

| **Model** | **MAE** | **MSE** | **RMSE** | **R2** |
| --- | --- | --- | --- | --- |
| **Assay CTX** | 1.729 ± 0.357 | 4.792 ± 1.711 | 2.162 ± 0.381 | -0.898 ± 0.665 |
| **ACL-DTI** | 0.744 ± 0.019 | 0.969 ± 0.047 | 0.984 ± 0.024 | 0.615 ± 0.016 |
| **MAMMAL-AsAP** | 0.795 ± 0.043 | 1.142 ± 0.120 | 1.067 ± 0.057 | 0.546 ± 0.048 |
| **Twinbooster** | $1.077 \pm 0.008$ | $1.872 \pm 0.020$ | $1.368 \pm 0.007$ | $0.258 \pm 0.011$ |

### Results of the Ablation Study

**Table 11:**
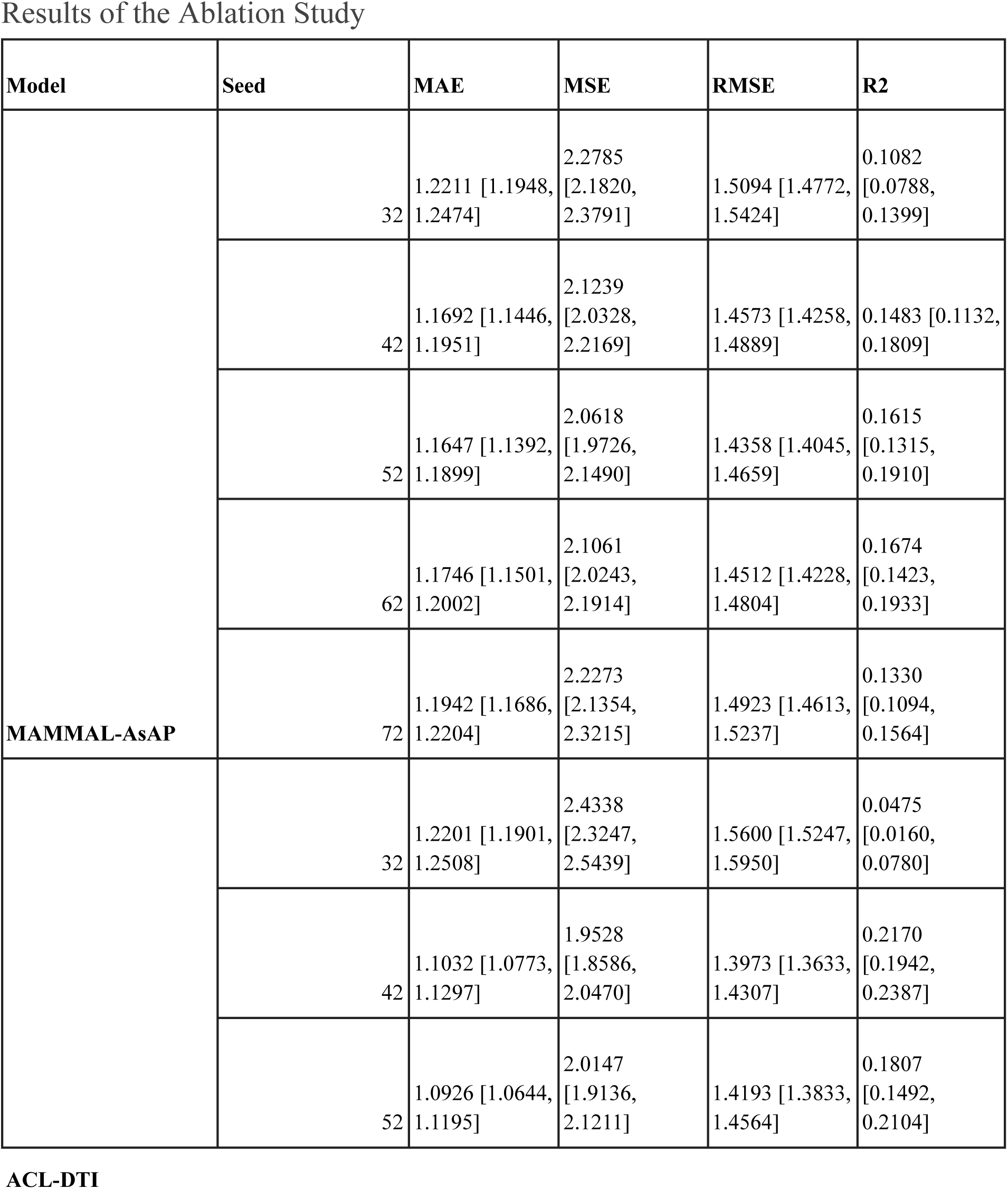
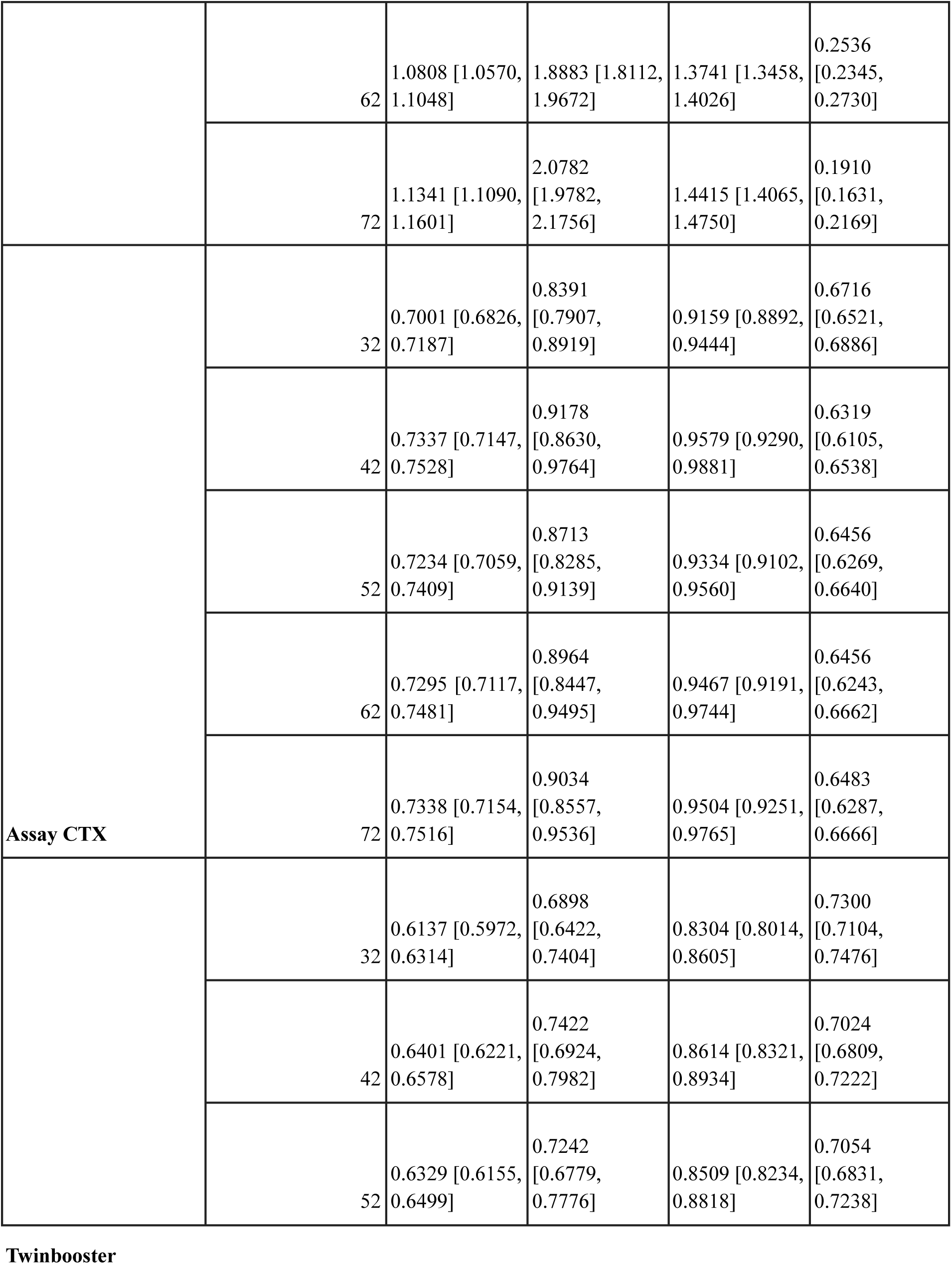
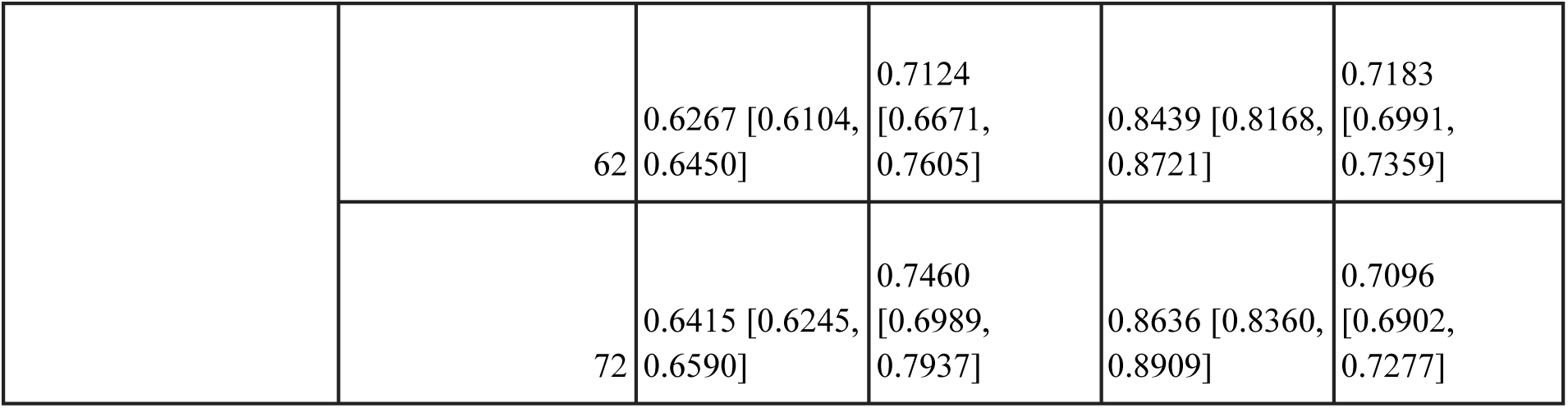
Ablation results for models using only drug sequence and assay description from the three input modalities.

**Table 12:** Average model performance using only drug sequence and assay description modalities across five random seeds. Values are reported as mean ± standard deviation across seeds.

| Model | MAE | MSE | RMSE | R2 |
| --- | --- | --- | --- | --- |
| Assay CTX | 0.724 ± 0.014 | 0.886 ± 0.031 | 0.941 ± 0.017 | 0.649 ± 0.014 |
| ACL-DTI | 1.126 ± 0.056 | 2.074 ± 0.213 | 1.438 ± 0.072 | 0.178 ± 0.078 |
| MAMMAL-AsAP | 1.185 ± 0.023 | 2.160 ± 0.090 | 1.469 ± 0.031 | 0.144 ± 0.024 |
| Twinbooster | 0.631 ± 0.011 | 0.723 ± 0.023 | 0.850 ± 0.014 | 0.713 ± 0.011 |

**Table 13:**
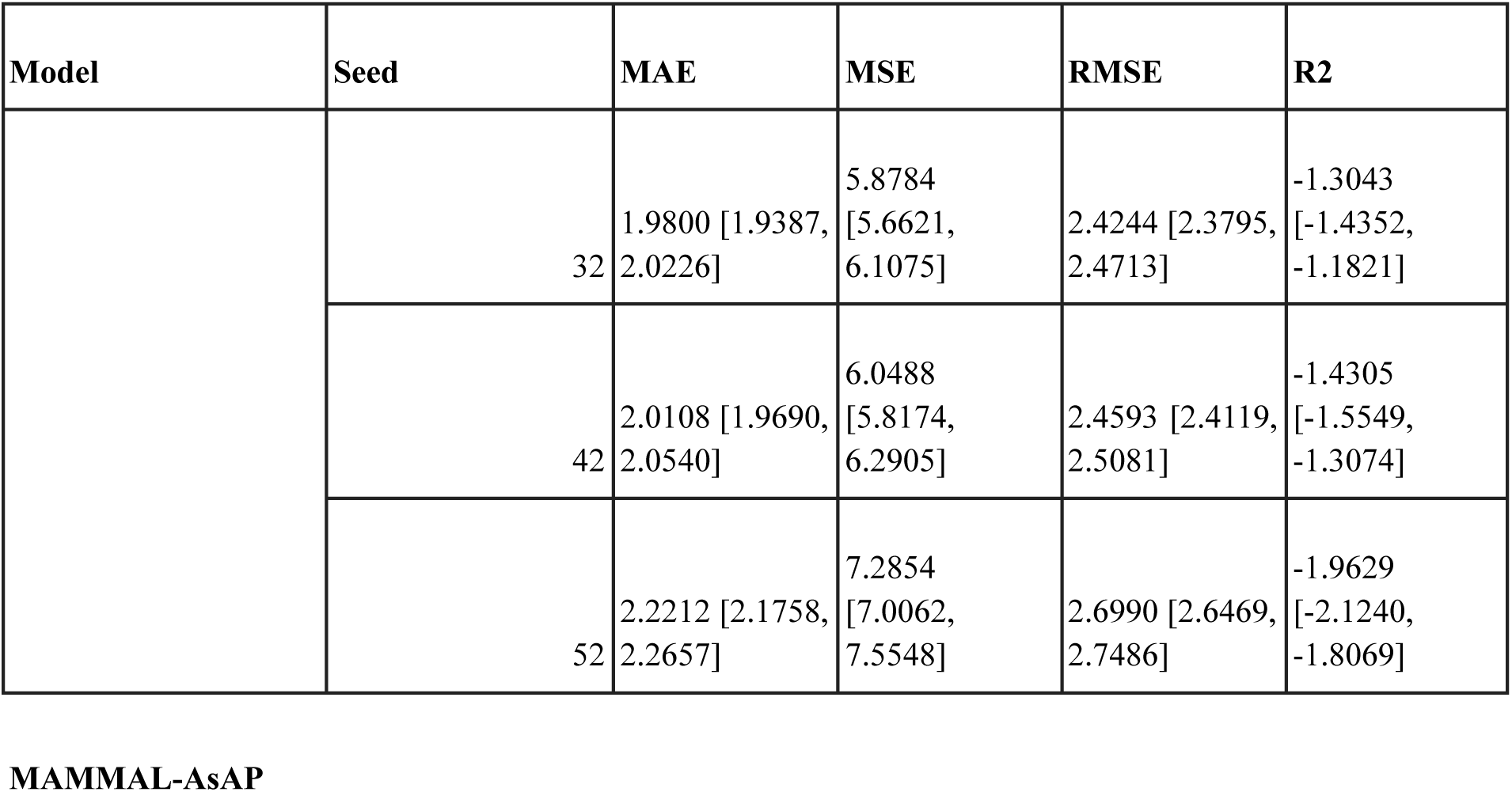
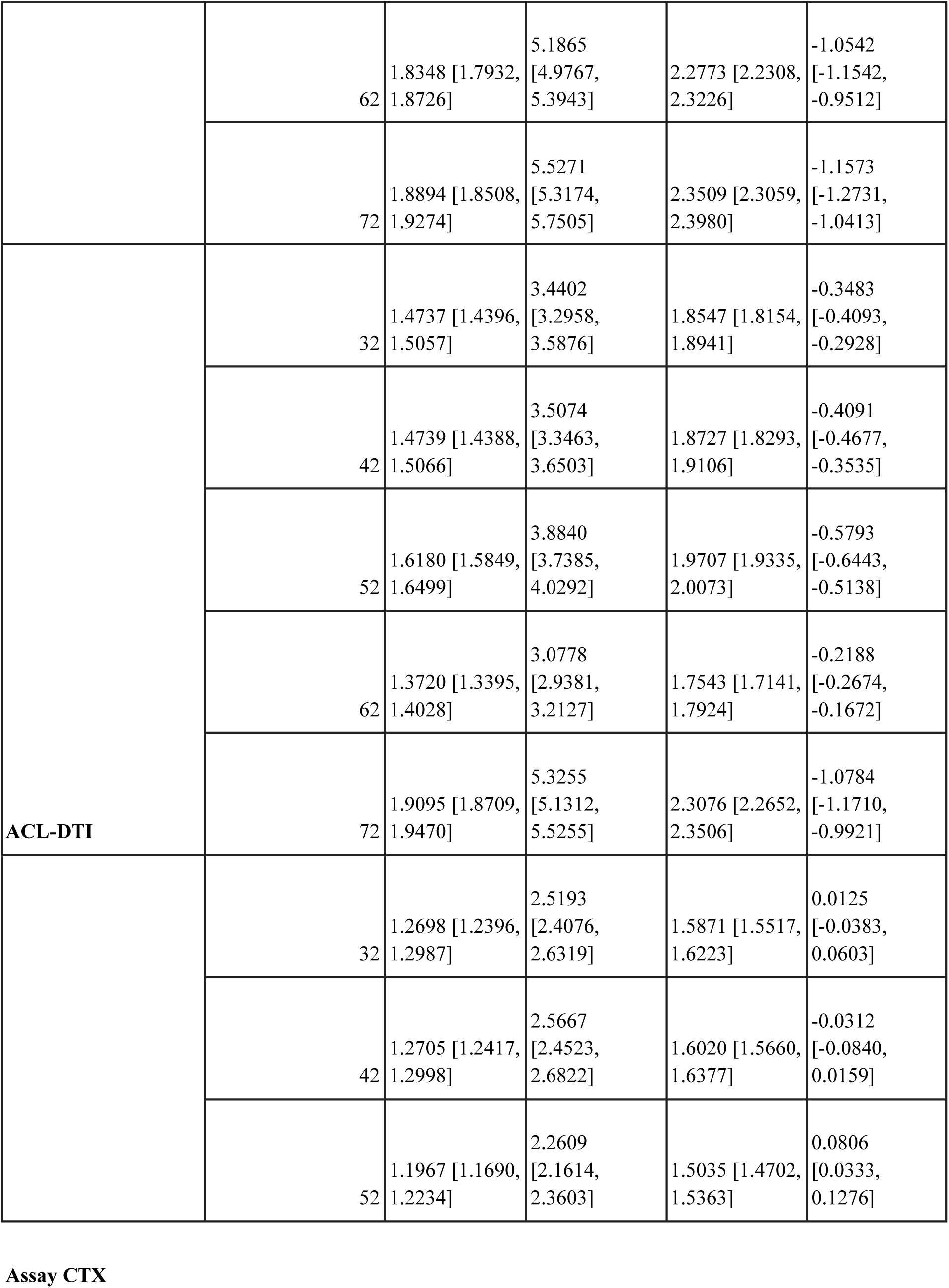
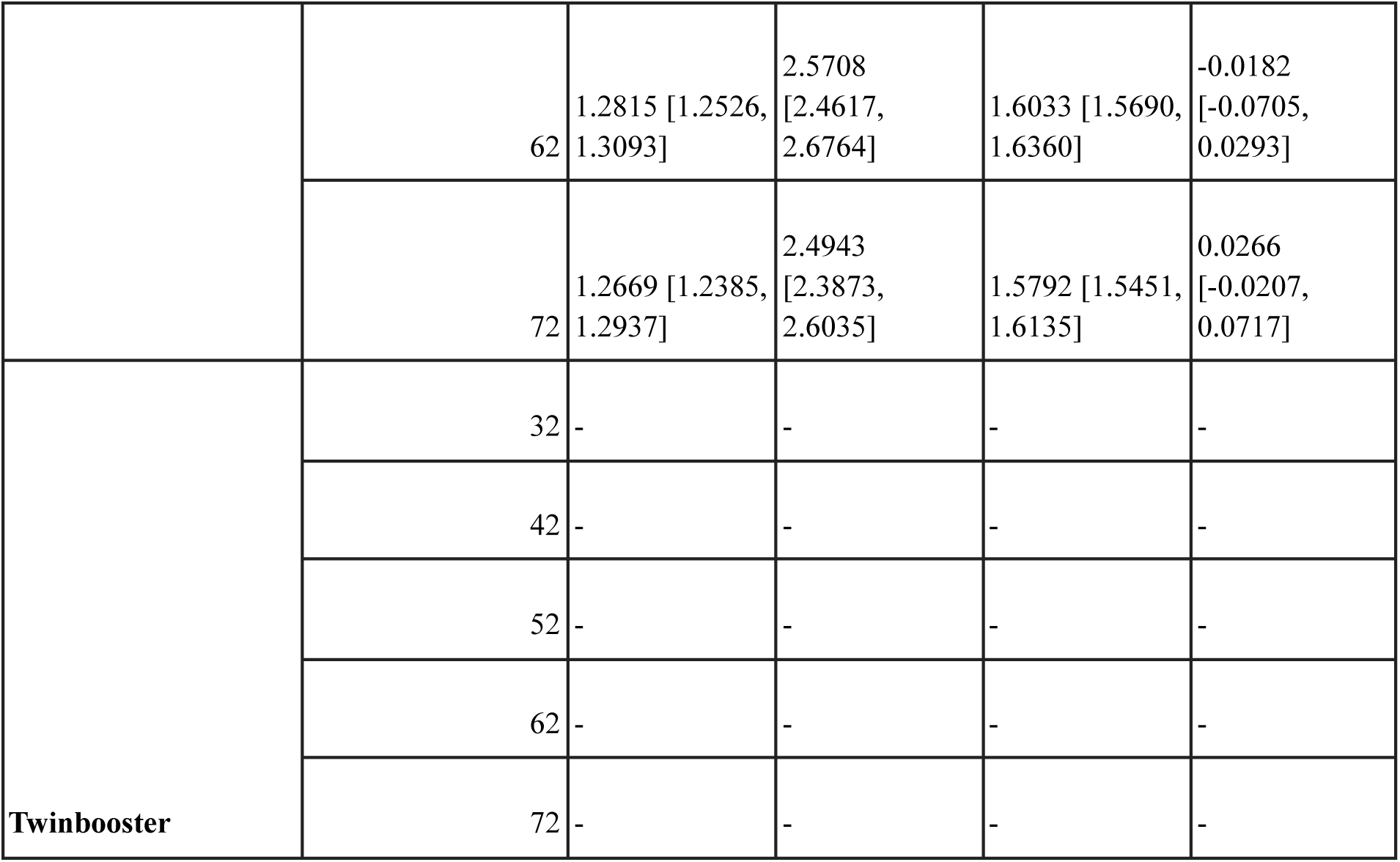
Ablation results for models using only target sequence and assay description from the three input modalities.

**Table 14:** Average model performance using only target sequence and assay description modalities across five random seeds. Values are reported as mean ± standard deviation across seeds.

| Model | MAE | MSE | RMSE | R2 |
| --- | --- | --- | --- | --- |
| Assay CTX | 1.257 ± 0.034 | 2.482 ± 0.128 | 1.575 ± 0.041 | 0.014 ± 0.044 |
| ACL-DTI | 1.569 ± 0.209 | 3.847 ± 0.875 | 1.952 ± 0.213 | -0.527 ± 0.335 |
| MAMMAL-AsAP | 1.987 ± 0.148 | 5.985 ± 0.799 | 2.442 ± 0.160 | -1.382 ± 0.355 |

**Table 15:**
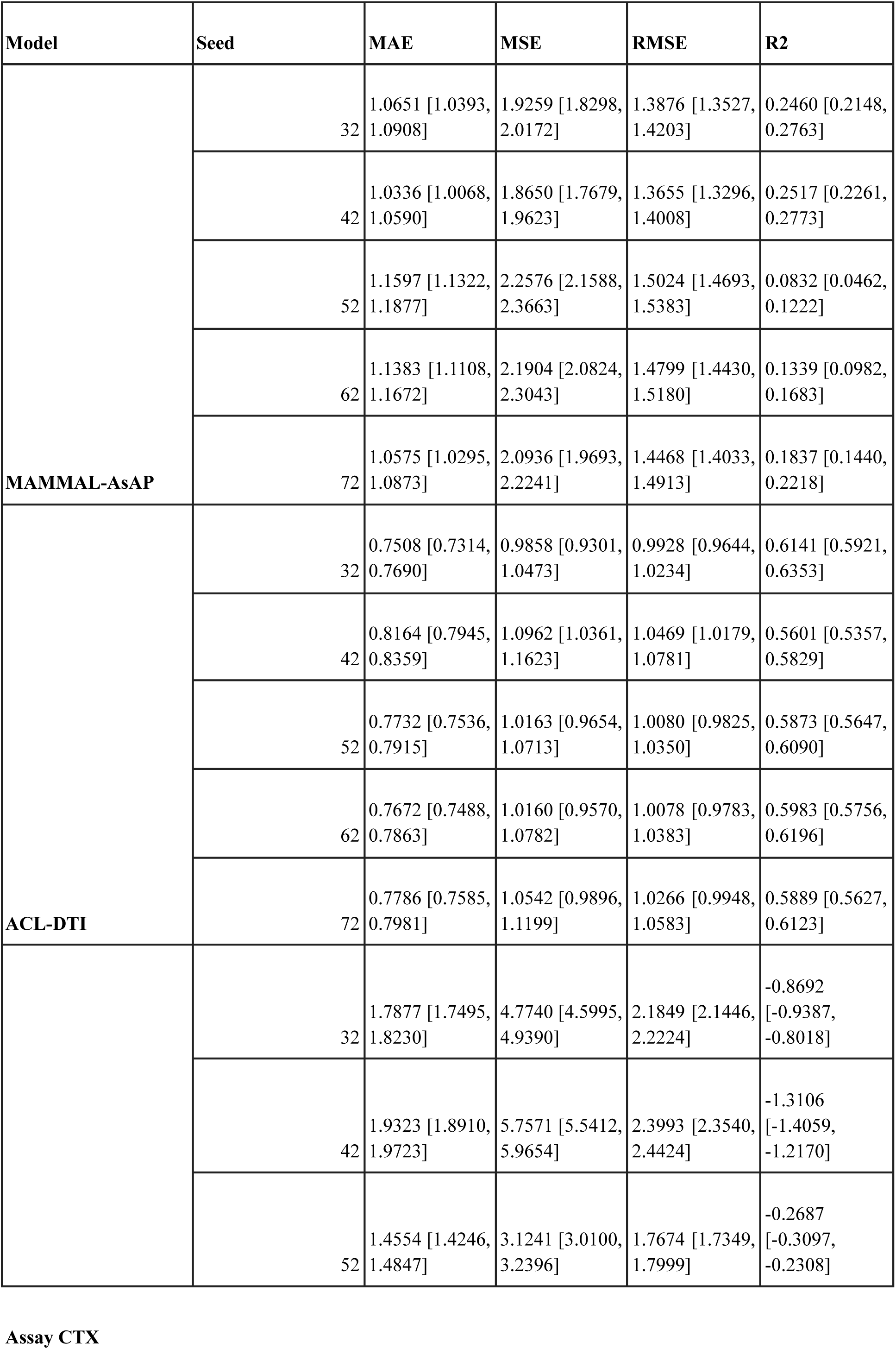
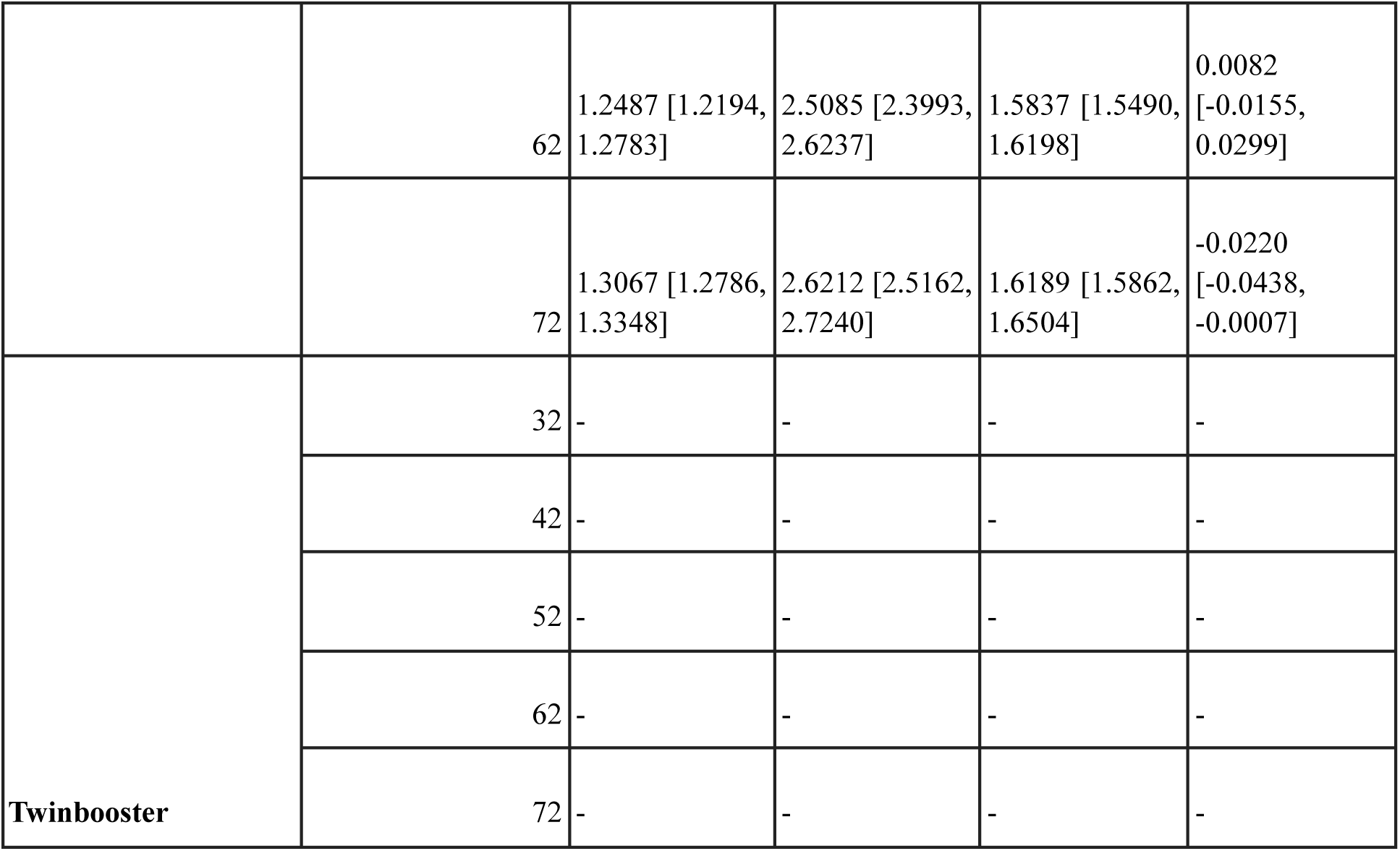
Ablation results for models using only drug sequence and target sequence from the three input modalities.

**Table 16:** Average model performance using only drug sequence and target sequence modalities across five random seeds. Values are reported as mean ± standard deviation across seeds.

| Model | MAE | MSE | RMSE | R2 |
| --- | --- | --- | --- | --- |
| Assay CTX | 1.546 ± 0.301 | 3.757 ± 1.439 | 1.911 ± 0.363 | -0.492 ± 0.577 |
| ACL-DTI | 0.777 ± 0.024 | 1.034 ± 0.043 | 1.016 ± 0.021 | 0.590 ± 0.020 |
| MAMMAL-AsAP | 1.091 ± 0.055 | 2.066 ± 0.168 | 1.436 ± 0.059 | 0.180 ± 0.072 |

**Table 17:**
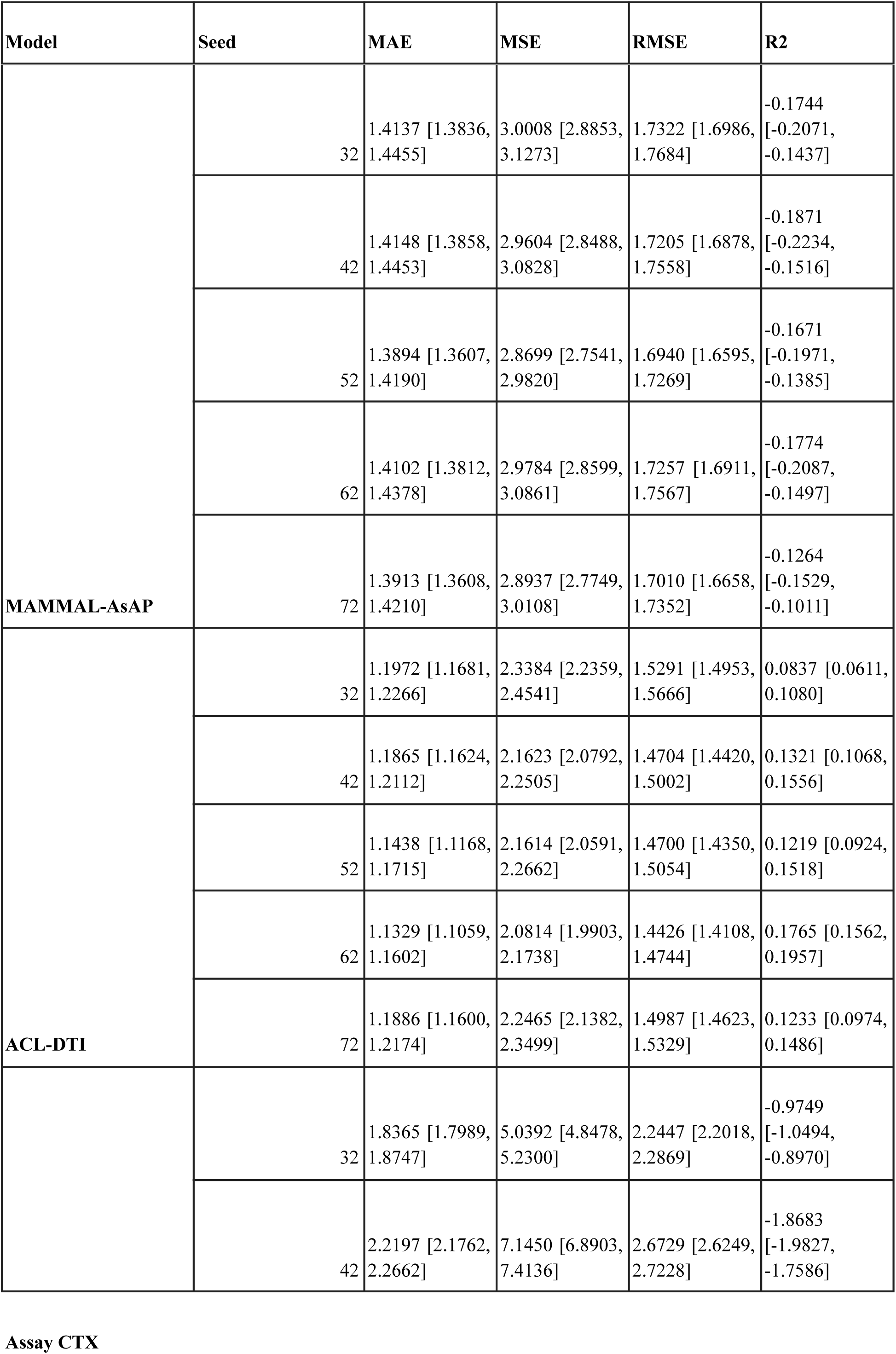
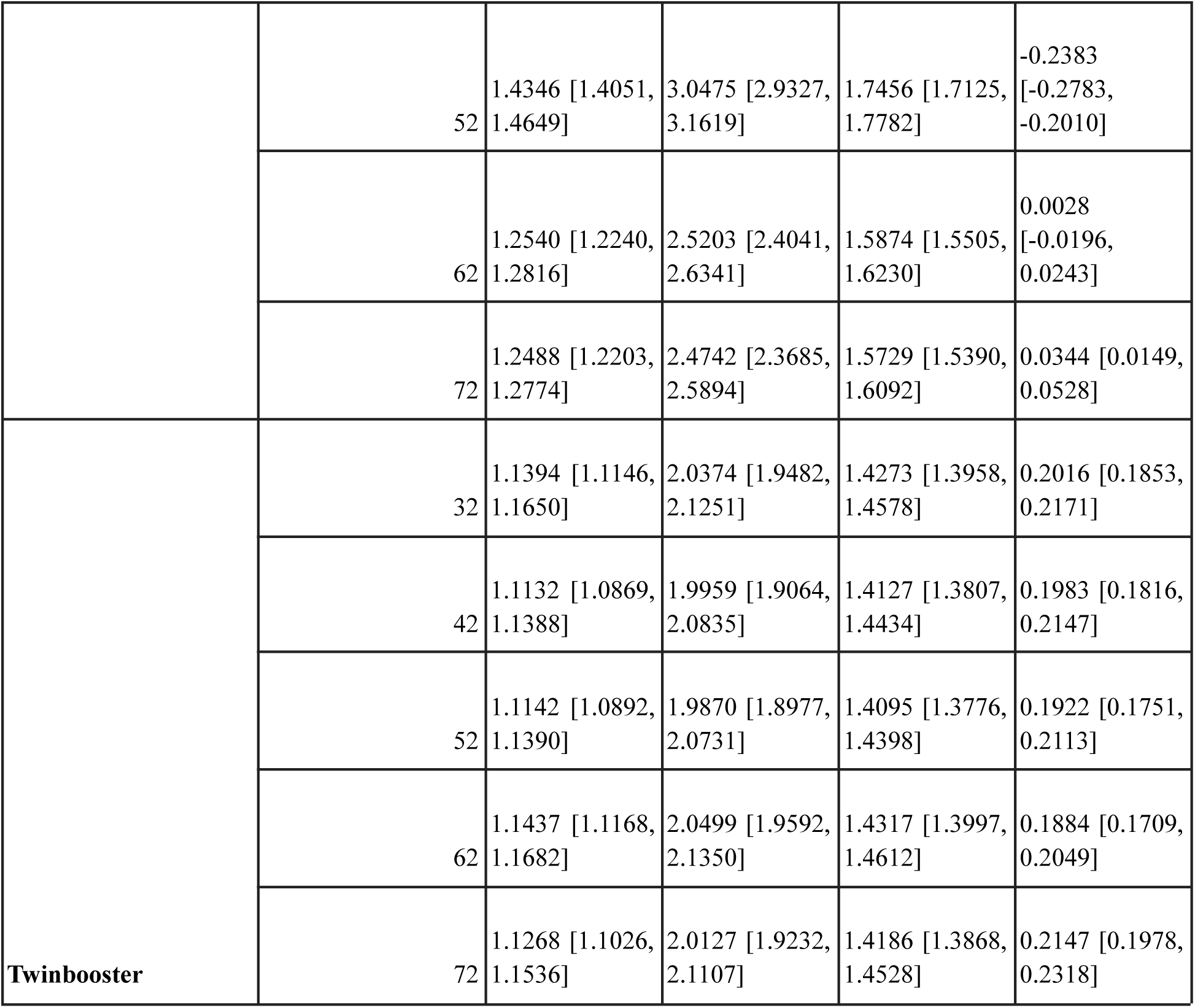
Ablation results for models using only drug sequence from the three input modalities.

**Table 18:** Average model performance using only drug sequence modality across five random seeds. Values are reported as mean ± standard deviation across seeds.

| **Model** | **MAE** | **MSE** | **RMSE** | **R2** |
| --- | --- | --- | --- | --- |
| **Assay CTX** | 1.599 ± 0.421 | 4.045 ± 2.024 | 1.965 ± 0.481 | -0.609 ± 0.813 |
| **ACL-DTI** | 1.170 ± 0.029 | 2.198 ± 0.098 | 1.482 ± 0.033 | 0.128 ± 0.033 |
| **MAMMAL-AsAP** | 1.404 ± 0.012 | 2.941 ± 0.056 | 1.715 ± 0.016 | -0.166 ± 0.024 |
| **Twinbooster** | 1.127 ± 0.014 | 2.017 ± 0.027 | 1.420 ± 0.009 | 0.199 ± 0.010 |

**Table 19:**
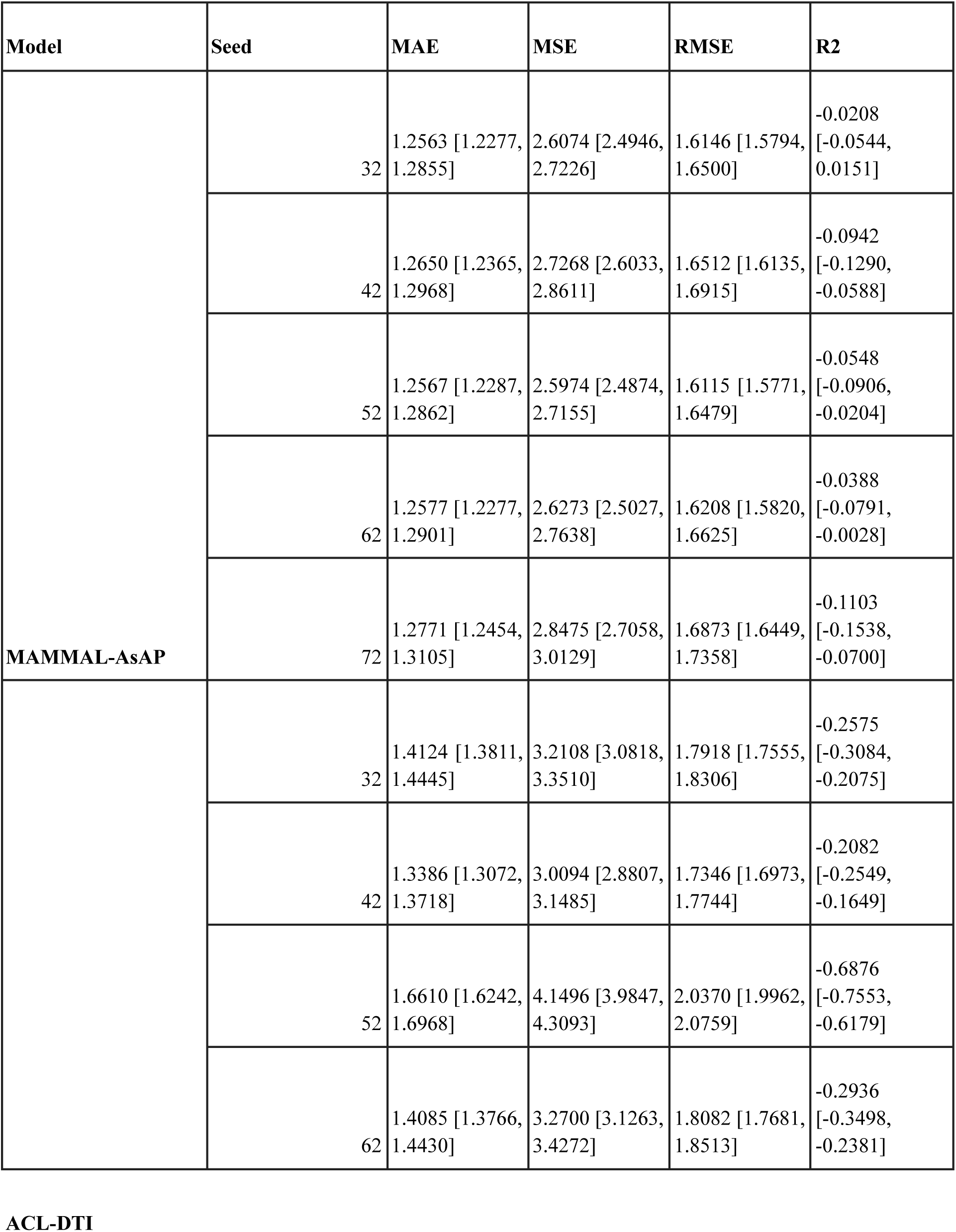
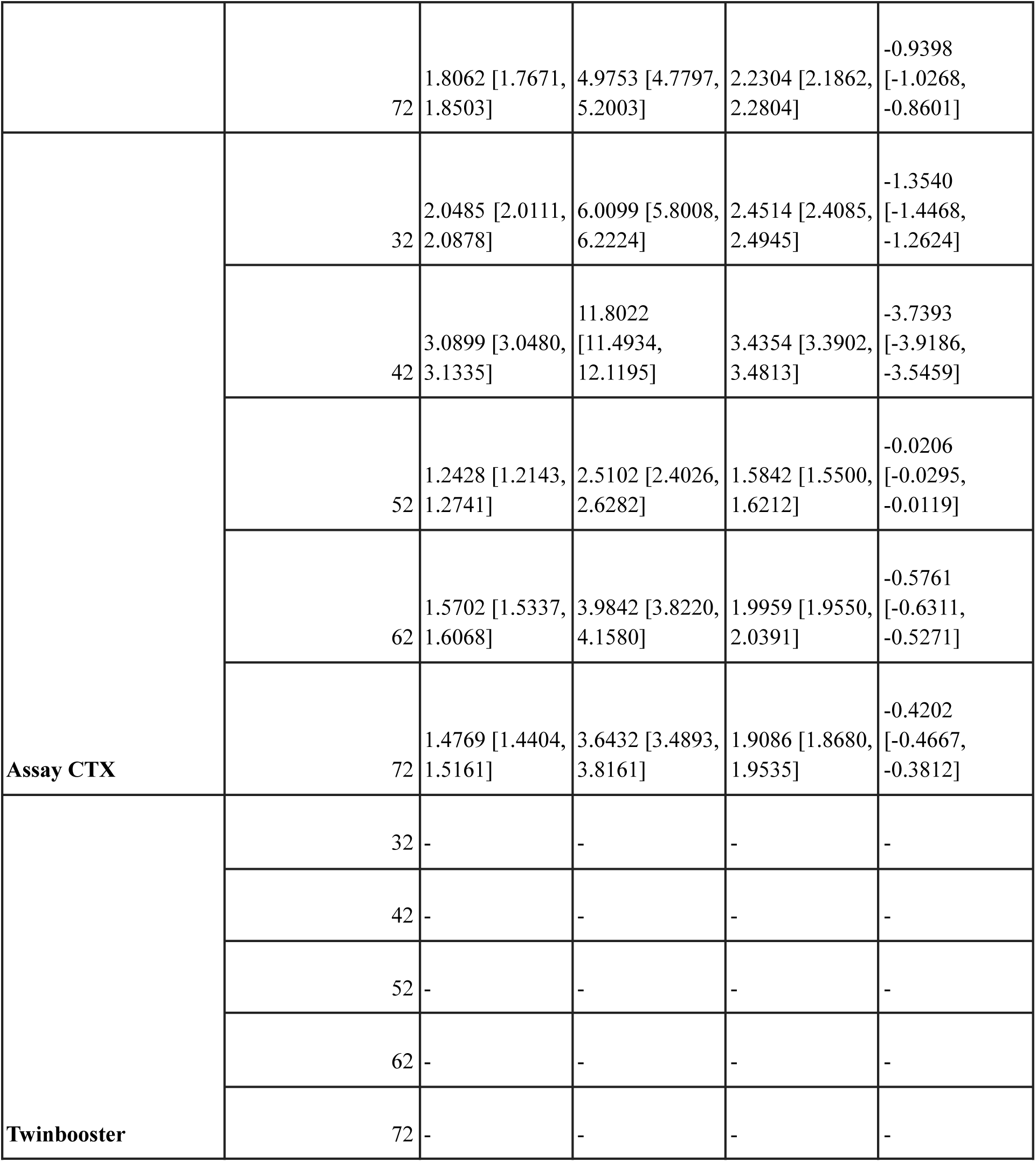
Ablation results for models using only target sequence from the three input modalities.

**Table 20:**
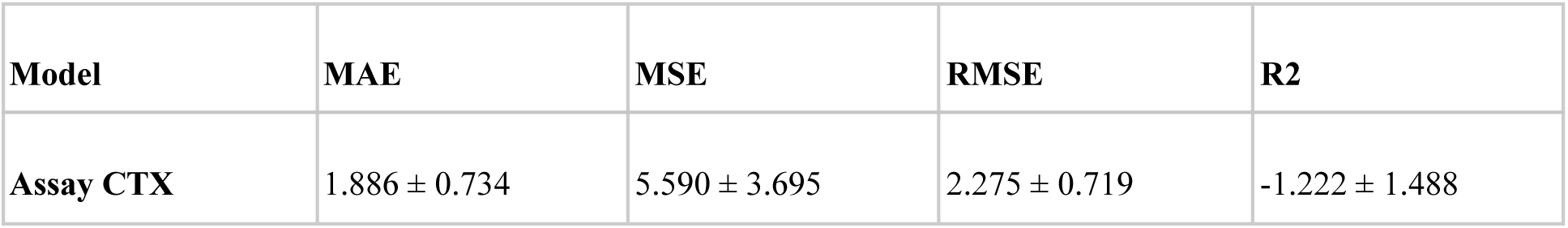
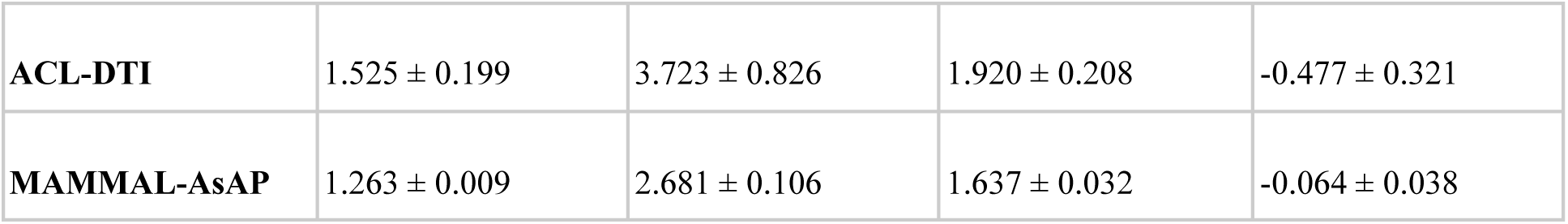
Average model performance using only target sequence modality across five random seeds. Values are reported as mean ± standard deviation across seeds.

| Model | MAE | MSE | RMSE | R2 |
| --- | --- | --- | --- | --- |
| Assay CTX | 1.886 ± 0.734 | 5.590 ± 3.695 | 2.275 ± 0.719 | -1.222 ± 1.488 |
| **ACL-DTI** | 1.525 ± 0.199 | 3.723 ± 0.826 | 1.920 ± 0.208 | -0.477 ± 0.321 |
| **MAMMAL-AsAP** | 1.263 ± 0.009 | 2.681 ± 0.106 | 1.637 ± 0.032 | -0.064 ± 0.038 |

**Table 21:**
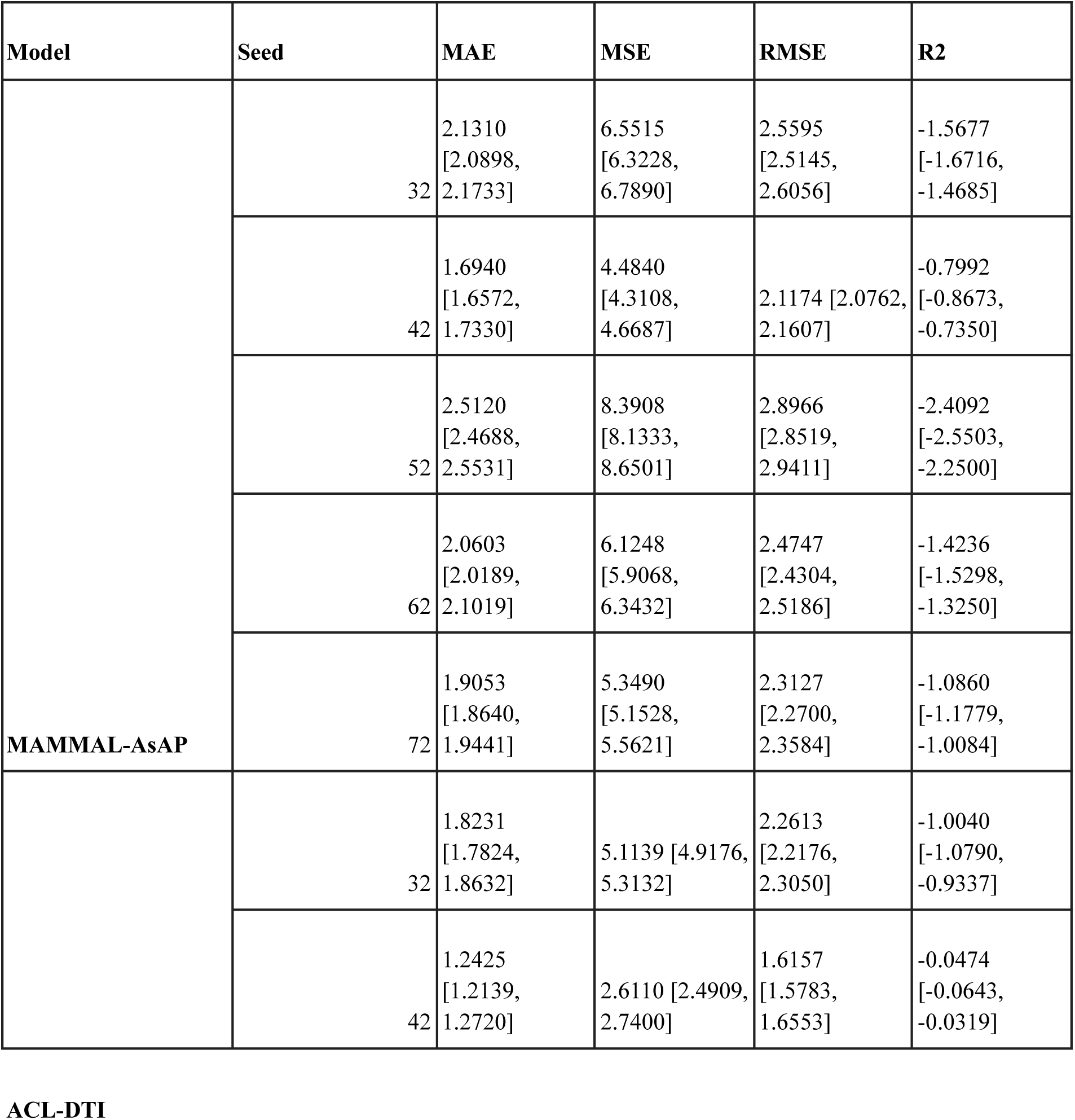
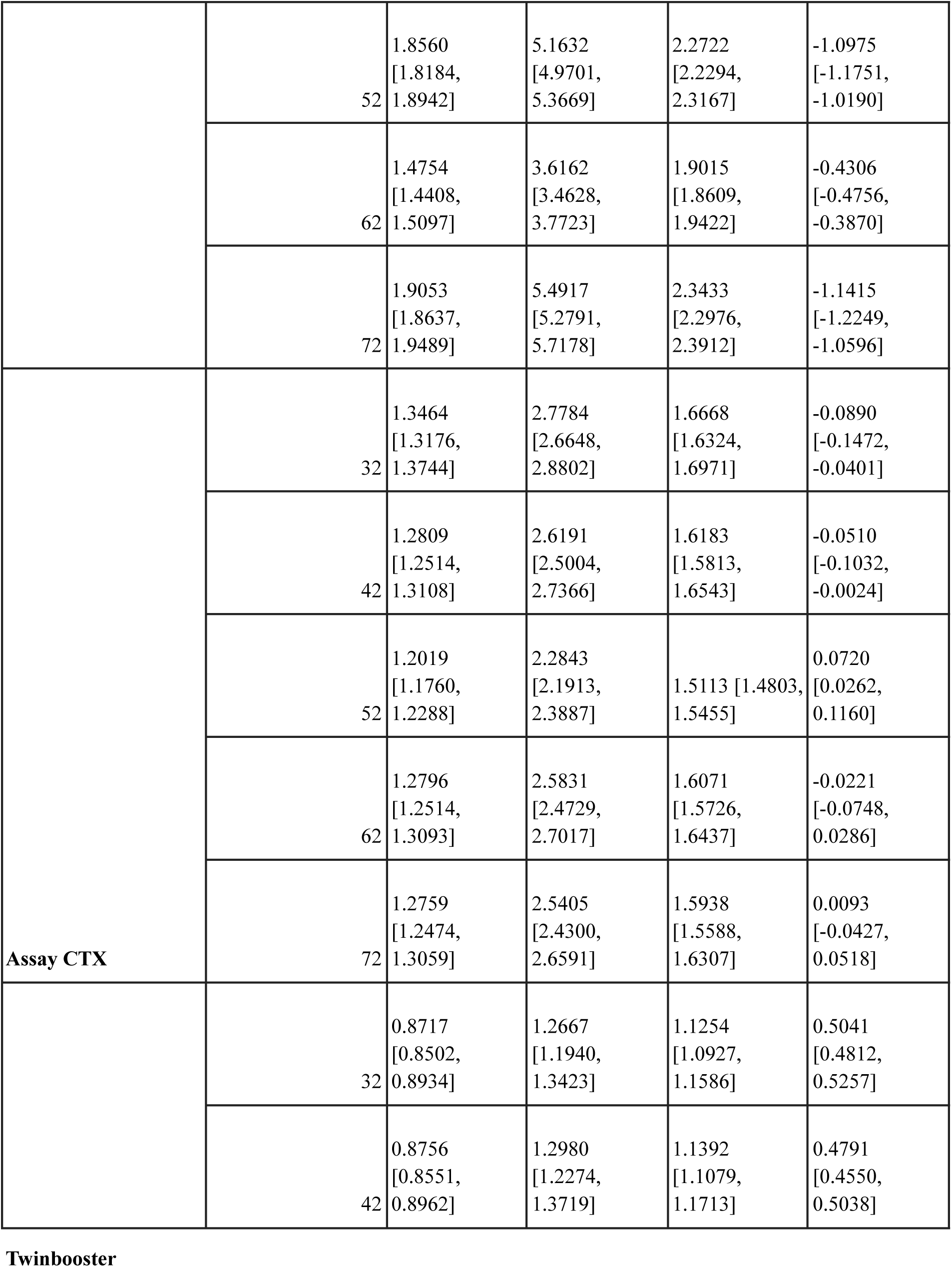
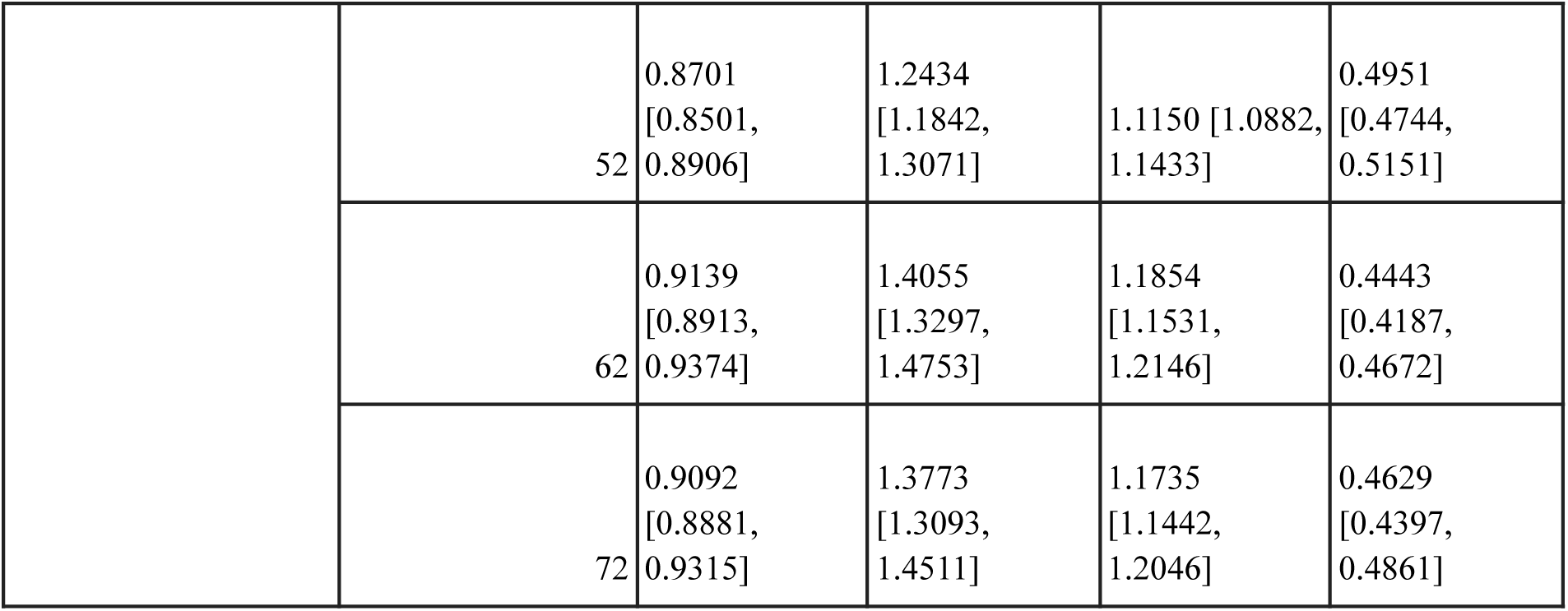
Ablation results for models using only assay description from the three input modalities.

**Table 22:** Average model performance using only assay description modality across five random seeds. Values are reported as mean ± standard deviation across seeds.

| Model | MAE | MSE | RMSE | R2 |
| --- | --- | --- | --- | --- |
| Assay CTX | 1.277 ± 0.051 | 2.561 ± 0.179 | 1.599 ± 0.056 | -0.016 ± 0.061 |
| ACL-DTI | 1.660 ± 0.289 | 4.399 ± 1.235 | 2.079 ± 0.311 | -0.744 ± 0.483 |
| MAMMAL-AsAP | 2.061 ± 0.303 | 6.180 ± 1.466 | 2.472 ± 0.291 | -1.457 ± 0.610 |
| Twinbooster | 0.888 ± 0.022 | 1.318 ± 0.070 | 1.148 ± 0.031 | 0.477 ± 0.024 |

